# Outcomes of early NIH-funded investigators: Experience of the National Institute of Allergy and Infectious Diseases

**DOI:** 10.1101/346049

**Authors:** Patricia A. Haggerty, Matthew J. Fenton

## Abstract

Survival of junior scientists in academic biomedical research is difficult in today’s highly competitive funding climate. National Institute of Health (NIH) data on first-time R01 grantees indicate the rate at which early investigators drop out from a NIH-supported research career is most rapid 4 to 5 years from the first R01 award. The factors associated with a high risk of dropping out, and whether these factors impact all junior investigators equally, are unclear. We identified a cohort of 1,496 investigators who received their first R01-equivalent (R01-e) awards from the National Institute of Allergy and Infectious Diseases between 2003 and 2010, and studied all their subsequent NIH grant applications through 2016. Ultimately, 57% of the cohort were successful in obtaining new R01-e funding, despite highly competitive conditions. Among those investigators who failed to compete successfully for new funding (43%), the average time to dropping out was 5 years. Investigators who successfully obtained new grants showed remarkable within-person consistency across multiple grant submission behaviors, including submitting more applications per year, more renewal applications, and more applications to multiple NIH Institutes. Funded investigators appeared to have two advantages over their unfunded peers at the outset: they had better scores on their first R01-e grants and they demonstrated an early ability to write applications that would be scored, not triaged. The cohort rapidly segregated into two very different groups on the basis of PI consistency in the quality and frequency of applications submitted after their first R01-e award. Lastly, we identified a number of specific demographic factors, intitutional characteristics, and grant submission behaviors that were associated with successful outcomes, and assessed their predictive value and relative importance for the likelihood of obtaining additional NIH funding.

## Introduction

Today, young scientists launching careers in biomedical research face a long, demanding path. The path includes years of post-graduate training, chronically low salaries, intense competition, historically low success rates for obtaining NIH funding, and a dearth of academic employment opportunities for independent scientists, given that the growth in number of advanced-degree graduates has outstripped the pace of research faculty positions opening [1-5]. Alberts et al. [6] attributed current systemic flaws in biomedical research in the United States (US) to a long-standing assumption that support and funding for this enterprise would expand almost indefinitely, a notion reinforced by the doubling of the NIH budget from 1999 to 2003. By the time the budget-doubling period ended, institutional expansion and growth of the scientific workforce resulted in a demand for research funds that far exceeded the availability of funds. Teitelbaum [7] described this disparity between supply and demand as the “structural disequilibrium” of research funding. This disparity was worsened by the US economic recession that began in 2008 and by the sequestration of the federal budget in 2013. As a result, NIH success rates declined to historic lows between 2003 and 2013 [8, 9], with little subsequent improvement.

Many in the field are concerned that new scientists will be discouraged from pursuing academic careers in the current climate. Stiff competition for research funds, low paylines, and poor job prospects are likely to drive talented investigators out of the biomedical workforce [4, 8, 10]. Even when new scientists secure an academic faculty position, their path to independence is still unsure, as evidenced by the continued increase in the average age of a NIH-funded investigator when obtainining their first R01 [11]). Moreover, NIH data (using cohorts from 1989, 1997, and 2003) show the rate of dropout (i.e. when an investigator fails to obtain a new or renewal R01-e grant award after the first one and stops applying) is greatest between 4 and 5 years from the first award [12]). Similar patterns were found using data from a cohort of NIAID first-time investigators from 1986 to 2003 (Fig 1). By 5 years, 68% of the NIAID cohort remained (32% dropped out), while 57% of the other NIH cohort remained (39% dropped out). The steep dropout between 4 and 5 years (red line in Fig 1) coincides with the duration of the first R01-e awards.

**Fig 1.**
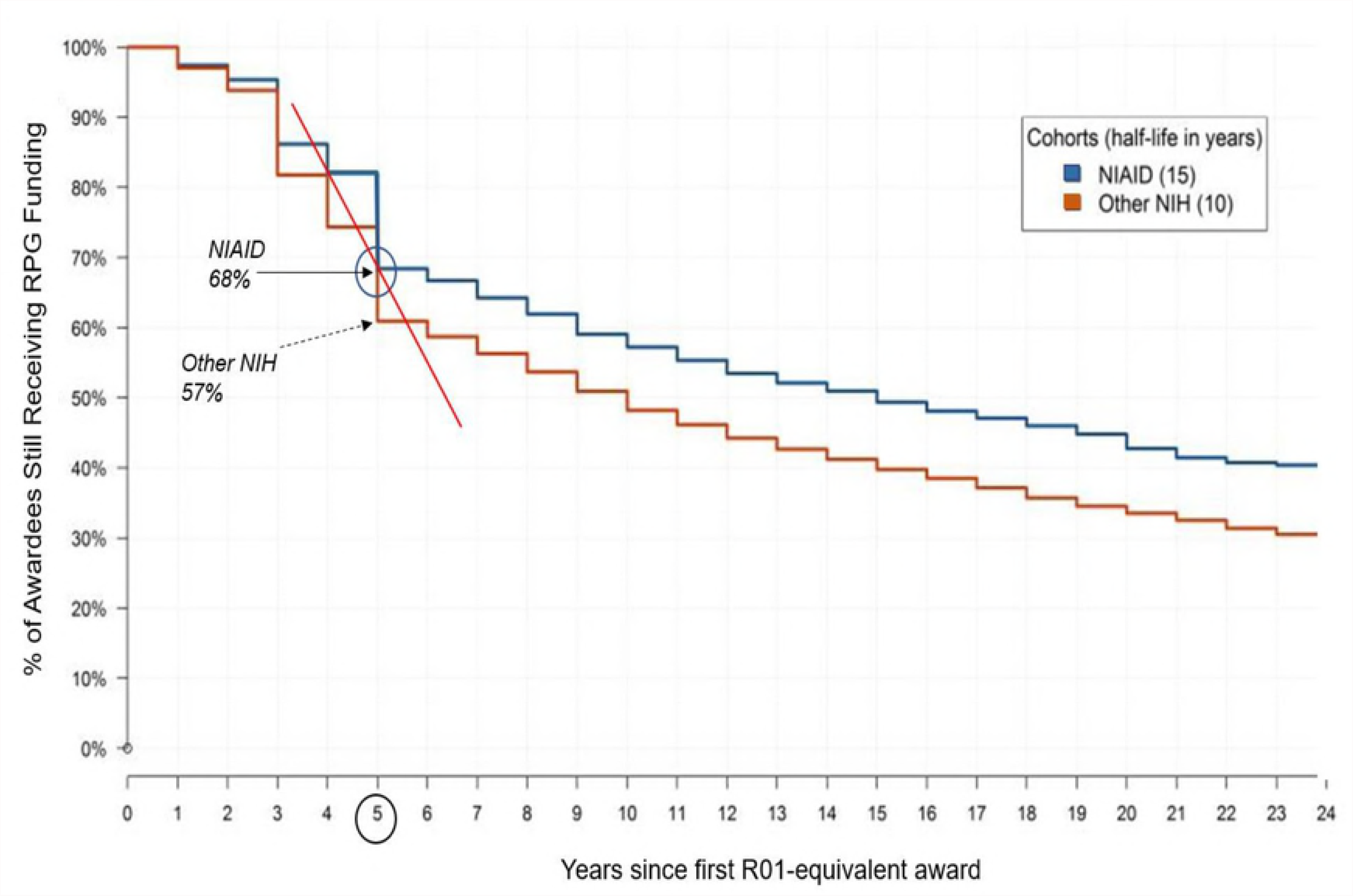
Length of Time Awardees Remain in NIH Applicant Pool After After the First R01-e Award. A Kaplan–Meier approach was used to measure the length of time investigators in each cohort remained in the NIH R01-e applicant pool after receiving their first R01-e awards. Y-axis: percent of investigators in each cohort who received an additional RPG award and remain in applicant pool. Investigators who do not remain in pool are considered to have ‘dropped out’. X-axis: years since receiving first R01-e award. Blue line: NIAID awardees. Orange line: other-NIH awardees. Solid red line: dropout slope between 4 and 5 years. Half of the NIAID cohort dropped out by 15 years after the first R01-e award (i.e. half-life 15 years); half of the other-NIH cohort dropped out by 10 years, or 50% sooner than the NIAID cohort.

What these prior reports did not address is whether there are specific risk factors leading to a high rate of dropout around the time that an investigator’s first R01 grant ends, and if so, if these factors impact all junior faculty equally. Furthermore, these prior studies did not discern whether there are characteristics and grant submission practices associated with investigators who are ultimately successful in obtaining future NIH funding, and those who are not. Armed with such knowledge, interventions might be developed to reduce the rate of dropout in this important pool of new scientists.

In order to better understand how first-time NIAID awardees compete for subsequent R01 awards, what their funding outcomes were, and when they drop out, we identified a cohort of principal investigators (PI) whose first R01-e awards were made by NIAID between between 2003 and 2010. We studied the cohort’s grant submissions and funding outcomes from the time of their first R01-e award through 2016. Our objectives were to learn: 1) what proportion of the cohort successfully competed for new or renewal NIH funding subsequent to their first award; 2) what were the grant funding outcomes and application submission behaviors of the PIs as they continued to apply for future funding; and 3) if there were demographic, institutional or other individual characteristics that differentiated successful and unsuccessful individuals.

## Methods

### Data Sources

All data used for this study came from the NIH database of information on extramural application and award records, known as Information for Management, Planning, Analysis, and Coordination II (IMPAC II). The NIH Query/View/Report (QVR) System was used to search the IMPAC II database and extract the data.

Personal demographic data on the cohort PIs, confidentially maintained by the NIH under the Privacy Act Systems of Record Notice 09-25-0036 [13] were provided to the authors by the NIH Office of Extramural Research, with permission. Regarding use of personally identifiable PI data, we followed the NIH policy stipulating: “All analyses conducted on date of birth, citizenship, gender, race, [and] ethnicity… data will report aggregate statistical findings only and will not identify individuals [14].

### First-time R01-Equivalent Awards

In addition to the R01, we include the following types of major research grants as R01-e: program projects and centers, cooperative agreements, other multi-project grants, and sub-projects of multi-project grants [15]. These grants are generally equivalent to the R01 in terms of cost, duration, effort, independence of the PI or Project Leader, and level of expertise required. NIH has historically considered a narrower range of grant types (referred to as activity codes) under the R01-e umbrella, but in programmatic contexts the activity codes considered to be R01-e have changed over time [16-17]. Unless otherwise specified, the term R01-e in this paper includes the broad range of grants mentioned above.

A small proportion of investigators, about 10%, received two first-time R01-e awards in the same fiscal year (FY). In these cases, we selected one of the two awards as their “first” award. To avoid confusion, we called the identified first award the “index award”, the application submitted for it the “index application”, and the FY the award was made in the “index fiscal year” (IFY).

### Study Time Frame

Our goal was to identify all PIs who received their index awards from NIAID between FYs 2003 and 2010, and to follow their subsequent grant submissions and outcomes. We chose 2003 as the cohort start year, because this was the end of the NIH budget doubling period [18]. We stopped the cohort at 2010 to allow sufficient time for first-time R01 awardees from this year to complete at least one 4- or 5-year project and apply for another.

The overall time frame of the study is from FYs 2003 through 2016. More precisely, for each investigator, the time frame is from the date of their index award until their final R01-e application, or through FY 2016, whichever came first. Thus, investigators who received their index awards in 2003, the first cohort year, were followed up to 13 years, investigators who received their index awards in 2004 were followed up to 12 years, and so on. Investigators who received index awards in the latest cohort year, 2010, were followed up to 6 years.

### Identification of Cohort PIs

From IMPAC II we extracted all competing R01-e awards made by NIAID between 1970 and 2010, excluding awards paid with funds appropriated under the American Recovery and Reinvestment Act of 2009. From this data set of 25,125 awards, we selected all awards made to PIs who were formerly NIH “New Investigators” prior to receiving that award [19]. Awards made to established investigators were omitted from the data set. More details of the steps we used to identify these awards and awardees are included S1 Appendix.

From the list of first-time awardees, 3 groups were distinguished: 1) those who received their first R01-e awards from NIAID; 2) those who received an R01-e award *other than R01* from NIAID and had not received an earlier R01 award from another NIH Institute (IC); and 3) those who received R01-e awards *other than R01* from NIAID *and* an earlier R01 award from another IC. We excluded the third group, because we wanted to focus on investigators who received their first awards from NIAID. Among the PIs who received R01-e awards *other than R01* from NIAID and no earlier R01 award (the second group), a very small number were subproject directors on a multi-project grant and these PI were kept in the cohort.

In total, we identified 1,496 investigators who received their first R01-e awards from NIAID between FYs 2003 and 2010. To distinguish these investigators from other established investigators, we called them “Early NIH-funded Investigators” (ENI).

### Application Data

In order to study the grant application submission behavior of the cohort, we took the unique PI identification number – the PI profile ID – of the 1,496 ENI and searched the IMPAC II database for all R01-e grant applications submitted by the cohort ENI to any NIH IC between 2003 and 2016.

In this study, we call every version of a grant application, whether it is the original version or one that was revised and resubmitted after an earlier unfunded version, an “application”. A new (NIH Type-1) application seeks funding for a new research project with different specific aims than any other project the PI has sought funding for. A renewal (NIH Type-2 or Type-9) application seeks an additional 4-5 years of funding for a research project that has already been funded by NIH for at least 4-5 years. Competitive supplement applications and applications withdrawn before peer review were not included.

The search extracted 12,964 applications, along with various project identifiers: PI identifiers, applicant institution information, the NIH IC assigned to the application, review information, and outcomes (funded or not funded). Eighteen percent (n = 2,365) of these applications were subprojects of multi-project grants.

### Application Outcomes

Examining the relationship between application outcomes and ENI funding success was an important part of this study. Here, we briefly describe how research project grant (RPG) application outcomes are determined at the NIH, and then discuss how we used cohort application outcome data.

Typically, during the NIH peer review process, about half of all RPG applications assigned to NIH study sections (committees) are “triaged”. That is, they are judged by the study section to be in the lower half, qualitatively, of all the applications assigned to the committee, and are designated as “noncompetitive”. Noncompetitive applications do not receive full discussion at the study section meeting, and their scores are not reported.

Applications that are not triaged receive a full discussion at the study section meeting and an overall numerical impact (or “priority”) score. Investigator-initiated R01 applications (i.e. most R01s) also receive a percentile score. The percentile score is based on a ranking of all the impact scores assigned by the committee in the previous 12 months. An application ranked in the 5^th^ percentile is considered more meritorious than 95% of the applications reviewed by that committee. Percentile scoring is intended to standardize impact scores across study sections that may have different scoring behaviors. R01 applications responding to a request for applications (RFA) and other R01-e applications are generally not percentiled.

NIAID establishes award thresholds from percentile ranks – called “paylines” – up to which nearly all R01 applications will be funded. For applications that are not percentiled, paylines are typically expressed as a priority score [20].

Therefore, the ENI applications included in this study had 3 possible outcomes: 1) triaged, unscored, not considered for funding; 2) scored, above the payline, ususally not funded; or 3) scored, within the payline, and funded. The majority of RPG applications that are not triaged are in the second category, i.e. initially judged to be competitive, but usually not funded. Many of these are subsequently revised and resubmitted for another round of peer review and funding consideration. Some applications that score above the payline may be funded under IC-specific funding rules.

Analysis of application outcomes was complicated by several factors: 1) in 2009, a new scoring system was introduced as part of the NIH Enhancing Peer Review initiative that changed scoring from a 0 to 500 point scale, to a 1 to 9 point scale [21]; 2) among the non-triaged applications, 20% had numerical priority scores but no percentile ranks; and 3) subproject applications (18% of all applications) had no triage identifiers, priority scores or percentile ranks.

For applications that were not triaged, the only valid metric for comparison purposes was the percentile rank. As noted, priority scores were subject to wide variation in study section behavior, so they could not be used. Therefore, for applications that had priority scores but no percentile ranks, we extrapolated percentiles in the following manner. For any given numerical priority score on a non-percentiled application, we took all percentiled applications with the same numerical score, calculated the average of their percentiles and assigned that percentile value to the non-percentiled application. This approach worked for applications before and after the change in the peer review scoring system and allowed us to include more of the applications in the data set.

There was no practical way to attach percentile scores to subproject applications, so these were excluded from any analyses that required application percentile data.

### PI-level Metrics

The primary outcome variable in this study is ENI success in obtaining additional new or renewal NIH R01-e funding after the IFY. ENI who obtained at least one additional award are called “funded” ENI, and those who did not obtain any additional awards are called “unfunded” ENI. For as long as an ENI continued to submit R01-e applications (or through FY 2016 at the latest), regardless of whether they were funded or unfunded, we followed their submission behaviors and application outcomes.

Because applications submitted by individual ENI reflect the application quality and submission behavior of that specific ENI, our analyses could not be based on comparisons between all applications from funded and unfunded ENI without potentially introducing bias. For example, multiple applications from the same ENI could artificially inflate or deflate summary metrics used to compare the two groups. Therefore, we concentrated on identifying comparisons at the person- (or PI-) level. We did use the application data to derive several PI-level metrics which we collectively called the PI SCORECARD. The values of items in the PI SCORECARD were based on the applications submitted by the ENI while s/he was in the study and included: SCORE (the average application score of all the ENI’s non-triaged applications); QUALITY (the proportion of all the ENI’s applications that were triaged, i.e. not scored at peer review, considered not competitive); FREQUENCY (the average number of applications submitted by the PI per year); SPEED (the length of time between the index award and first subsequent grant submission); REACH (the proportion of the PI’s applications submitted to a single NIH IC (versus to multiple ICs)); RENEW (the proportion of the PI’s applications that were renewal applications); RESUB (the proportion of the PI’s applications that were resubmissions, i.e. previously peer reviewed but not funded, formerly called amended applications); ACTIVE (the length of time the PI remained in the R01-e applicant pool); and INDEX (the PI’s index award percentile score).

### Project Start Dates

A critical application-associated data field in this study was the project start date. Project start date was essential for: 1) identifying ENI index awards; 2) chronologically ordering applications for each ENI; and 3) calculating the time between the index award and the ENI’s subsequent applications.

For many applications in our data set, project start dates were missing or inaccurate, due to a variety of reasons. (For more information about project start date, see S1 Appendix.) Therefore, we chose to use a different parameter altogether as a proxy for project start date. All but 57 applications had Council Dates (i.e. the meeting date of the National Advisory Allergy and Infectious Diseases Council). We took the NIAID average time from Council Date to notice-of-award date (4 months) and added this to the application Council Date to derive an estimated project start date. We applied this approach uniformly to all applications, except for the 57 without Council Dates. Fortunately, the latter had accurate project start dates, which we used.

### Statistical Methods

For comparisons between funded and unfunded ENI according to independent categorical variables, we used Pearson’s χ^2^ test. For comparisons between the two groups according to independent continuous variables, we used the Welch Two Sample t-Test. The full cohort of ENI (n=1,496) was used in comparisons between funded and unfunded ENI according to the inherent independent variables, i.e. demographic, institutional, and PI background characteristics. In contrast, comparisons between the two groups according to PI SCORECARD items were limited to just the ENI who submitted additional grant applications after the IFY (n = 1,322, or 88% of the cohort). Our rationale was that the strength of associations of these variables with funding outcome may have been influenced as a consequence of repeated grant writing.

To analyze the effects of independent (predictor) variables on the likelihood of ENI funding, we used univariate and multivariate logistic regression analyses. To identify the relative importance of each of the variables in predicting ENI funding success when all variables were included in a multivariate model, we used random forests (RF), a machine learning alorithm that evaluates the importance of variables by estimating the change (i.e. the prediction error) in a model quality score that occurs when any single variable is randomly permuted, while others are left unchanged [22]. Larger values of importance indicate stronger predictors, and values close to zero suggest the variable is not a good predictor. RF are popular because of their ability to deal with large numbers of covariates, non-linear associations, complex interactions and correlations between variables; RF have been used in many biomedical research fields [23-27]. In our RF variable importance (RFVI) analysis we converted all predictor variables into binomials, to avoid reported possible bias of RF when used with categorical variables with multiple levels, or correlated predictors [28, 29].

All statistical analyses were performed using R version 3.4.3, with packages plyr, dplyr, ggplot2, readxl, lmtest, and randomForest (Breiman and Cutler, 2018) [30-36]. Microsoft Excel 2016 was used for early conditioning of raw data extracted from IMPAC II.

## Results

### Cohort Descriptive Characteristics

An average of 13% of the ENI came from each of the 8 cohort years (2003 – 2010) (Table 2). Slightly more than half (52%) of the cohort came from the first 4 years (2003 – 2006), and 2003 had the largest number of ENI compared to all the other years.

**Table 1.** PI SCORECARD: Grant Submission Behaviors and Grant Quality Indices.

| PI Factor | Definition | Meaning of Factor Value |
| --- | --- | --- |
| **SCORE** | PI average application score (percentile) | Lower = stronger |
| **QUALITY** | % of PI's applications triaged | Lower = stronger |
| **INDEX** | Index award score (percentile) | Lower = stronger |
| **FREQUENCY** | PI's average number of applications per year | Higher = more |
| **SPEED** | Time (y) between index award and first subsequent R01-e application | Lower = faster |
| **REACH** | % of PI's applications sent to single NIH IC | Lower = more sent to other ICs |
| **RENEW** | % of PI's applications as renewals | Higher = more |
| **RESUB** | % of PI's applications as resubmissions | Higher = more |
| **ACTIVE** | Number of years PI remained in R01-e NIH applicant pool | Higher = longer |

**Table 2.** Number of ENI per Cohort Year and Percent of Total Cohort.

| Cohort Year* | # of ENI | % of Total Cohort |
| --- | --- | --- |
| 2003 | 236 | 16% |
| 2004 | 185 | 12% |
| 2005 | 194 | 13% |
| 2006 | 167 | 11% |
| 2007 | 199 | 13% |
| 2008 | 151 | 10% |
| 2009 | 197 | 13% |
| 2010 | 167 | 11% |
| Total Cohort | 1496 | 100% |

\*Cohort Year = FY in which ENI received index award and entered study cohort

### ENI Demographic Characteristics

Demographic characteristics of the cohort are shown in Table 3. Just under three quarters (73%) of the ENI were male. Of 1,370 ENI with known date of birth, the median age at receipt of the index award was 41.2 y (mean 42.6 y). Of 1,301 ENI with known birth countries, 75% were born in the US, and 25% in 66 other countries. The proportions of investigators by gender, birth country, and age at index award, are similar to overall NIH data [37, 38].

**Table 3.**
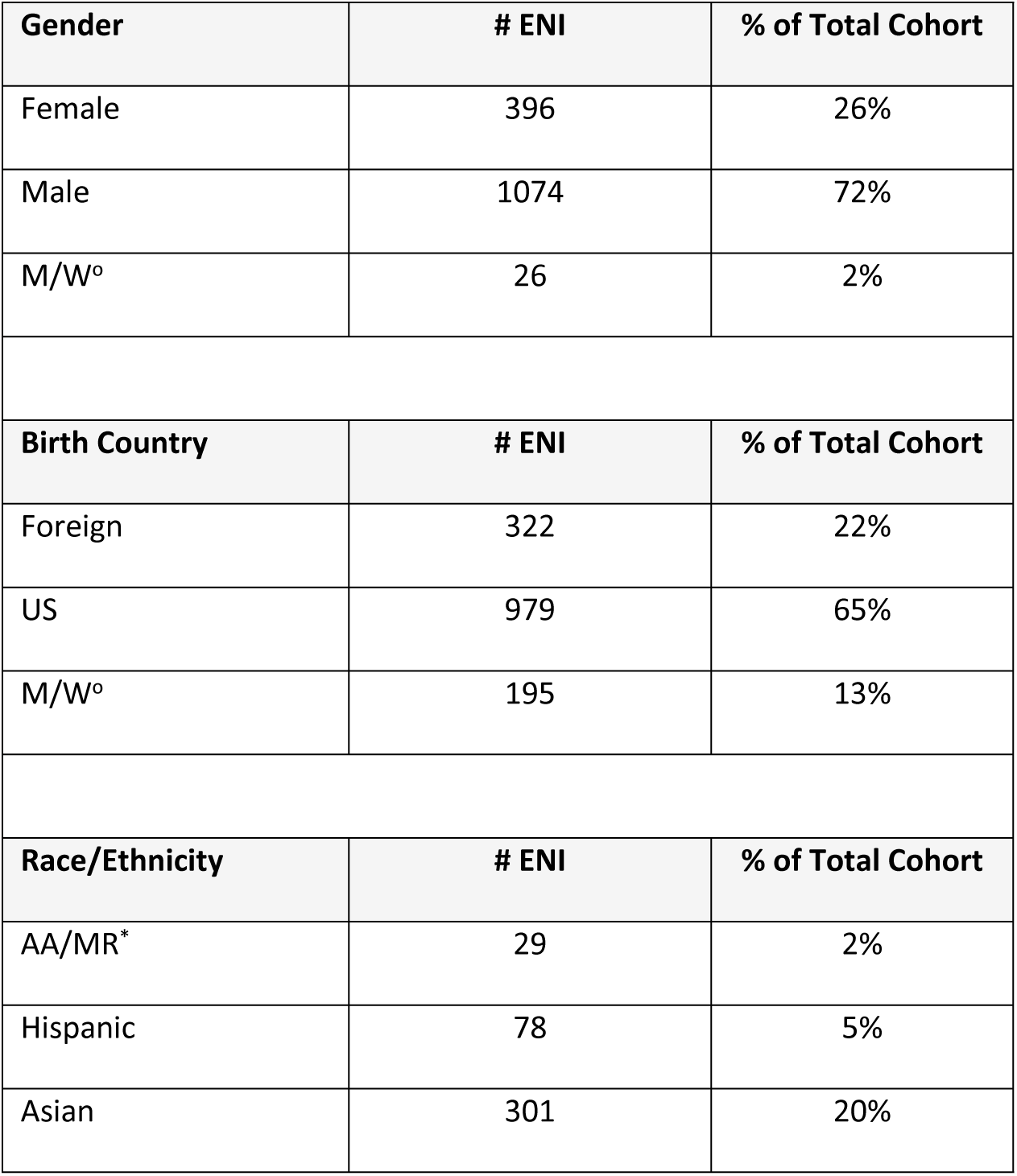
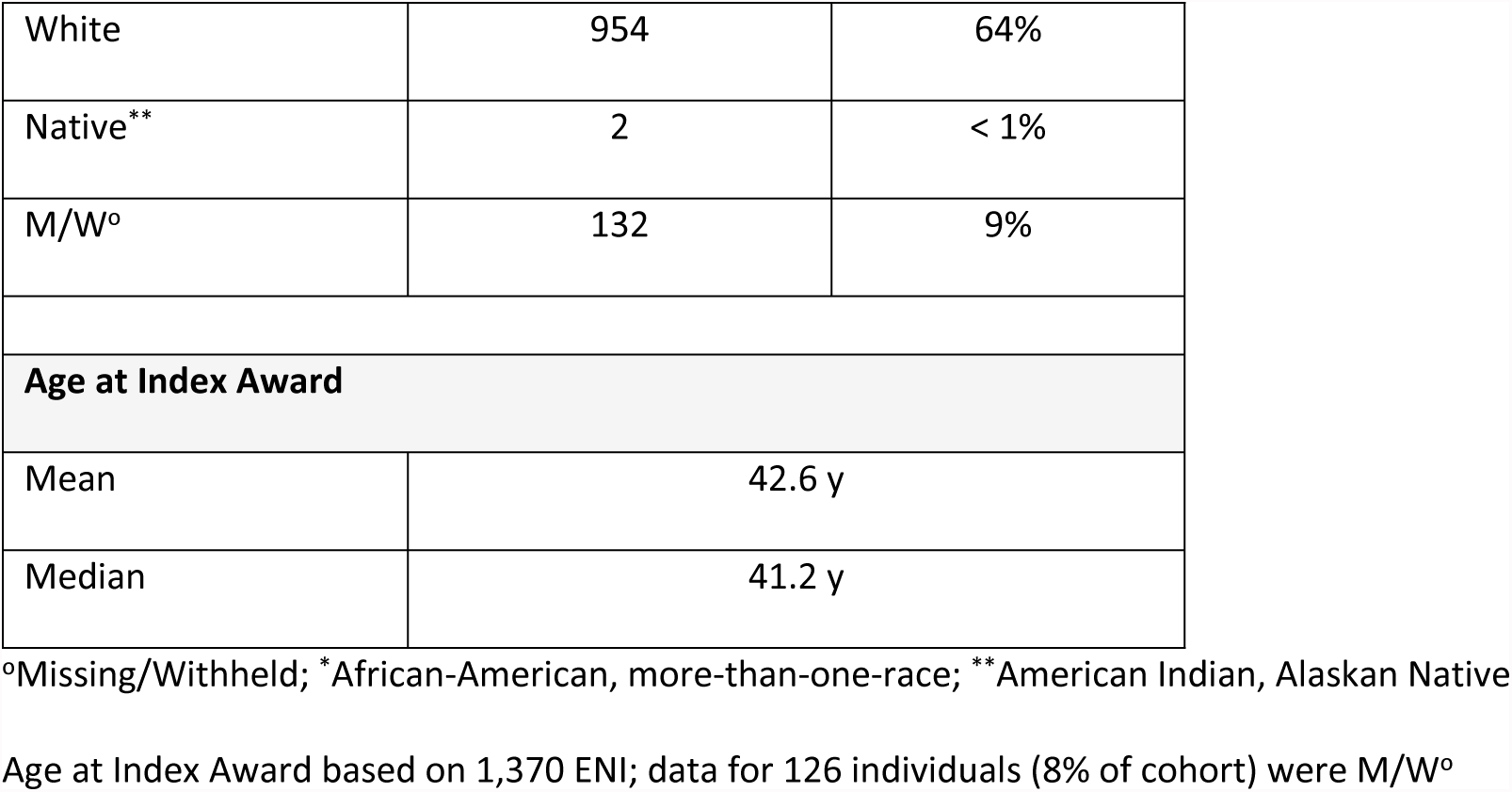
ENI Demographic Characteristics.

In terms of self-reported race and ethnicity, 64% of the ENI were white, 20% Asian, 5% Hispanic, 1.5% African American (AA), less than 1% more-than-one-race (MR), and less than 1% Native (American Indian or Alaskan Native). We combined the 22 AA and 9 MR ENI into a single group (AA/MR), representing 2% of the cohort. Compared to the NIH overall, the NIAID cohort had slightly higher representations of AA and Hispanic investigators, and a lower representation of white investigators. Between 1999 and 2012 the NIH had, on average, 1.2% Black, 3.5% Hispanic, and 79% White R01 awardees [39].

### ENI Background and Index Institution Characteristics

ENI terminal research or post-graduate clinical training degrees were categorized into 4 groups: MD, MD/PhD, PhD (or equivalent), and Other (Table 4). Almost 70% of ENI had PhD or equivalent degrees, 30% had MD or MD/PhD degrees, and 1% had other degrees. Seventeen percent of ENI were prior recipients of an NIH career development (i.e. “K”) award. All but 8% of ENI were employed at US institutions when they received their index award, and most institutions were non-medical school institutions of higher education (43%) or medical schools (29%). This distribution of ENI across these institution types parallels the historic distribution of institution types according to allocation of NIH grant funding [40].

**Table 4.** ENI Background and Index Institution Characteristics.

| Degree Group | # ENI | % of Total Cohort |
| --- | --- | --- |
| MD | 289 | 19% |
| MD, PhD | 172 | 11% |
| PhD (or equiv.) | 1041 | 68% |
| Other | 20 | 1% |
| **Prior K Award** | **# ENI** | **% of Total Cohort** |
| No | 1249 | 83% |
| Yes | 247 | 17% |
| **Institution US or Foreign** | **# ENI** | **% of Total Cohort** |
| Foreign | 125 | 8% |
| US | 1371 | 92% |
| **Institution Type** | **# ENI** | **% of Total Cohort** |
| Independent Hospital | 130 | 9% |
| Higher Education (non-medical school) | 636 | 43% |
| Other Health, Health-rel., Community Srvc. | 36 | 2% |
| Independent Research | 156 | 10% |
| Medical School | 429 | 29% |
| Company | 108 | 7% |
| Foreign | 1 | < 1% |

In this study, the term “institution” refers to the institution where the PI was employed at the time of receiving his/her index award. To characterize institutions further, we took the ENI from US, non-commercial institutions, and divided them into 3 roughly equal groups – or “tertiles” – according to the number of ENI employed at those institutions when they received their first R01-e grant (Table 5). There were 1,272 ENI, from 269 institutions, in this analysis; 181 ENI from 103 foreign institutions and 78 US companies were excluded. The first tertile included 406 ENI from institutions with 14 to 37 ENI per institution, or “high-ENI density” institutions. The second tertile included 454 ENI from institutions with 6 to 13 ENI per institution, or “medium-ENI density” institutions. The third tertile included 412 ENI from “low-ENI density” institutions, or institutions with 1 to 5 ENI per institution. Interestingly, just 75 of the 269 institutions (28%) were in the top two tertiles, while 194 institutions (72%) were in the third tertile. The high- and medium-ENI density institutions historically have been, and remain, among the NIH top-funded institutions [41].

**Table 5.**
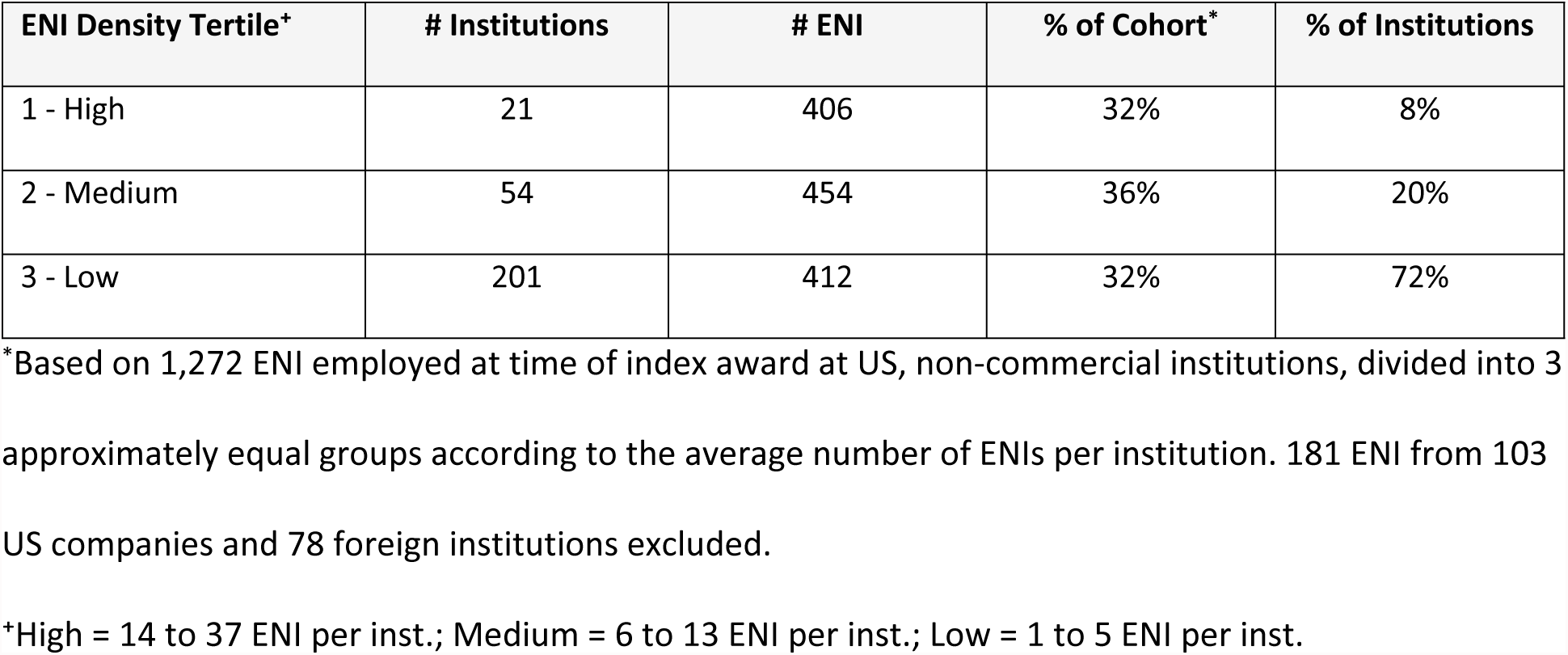
Institution ENI Density.

### Index Award Characteristics

ENI index awards included research projects, multi-project programs and centers, and multi-project cooperative agreements. More than three fourths of the awards were research project R01, 14% were research project U01, and the remaining 8% were multi-project awards. The distribution of index awards by activity type codes is shown in Fig 2.

**Fig 2.**
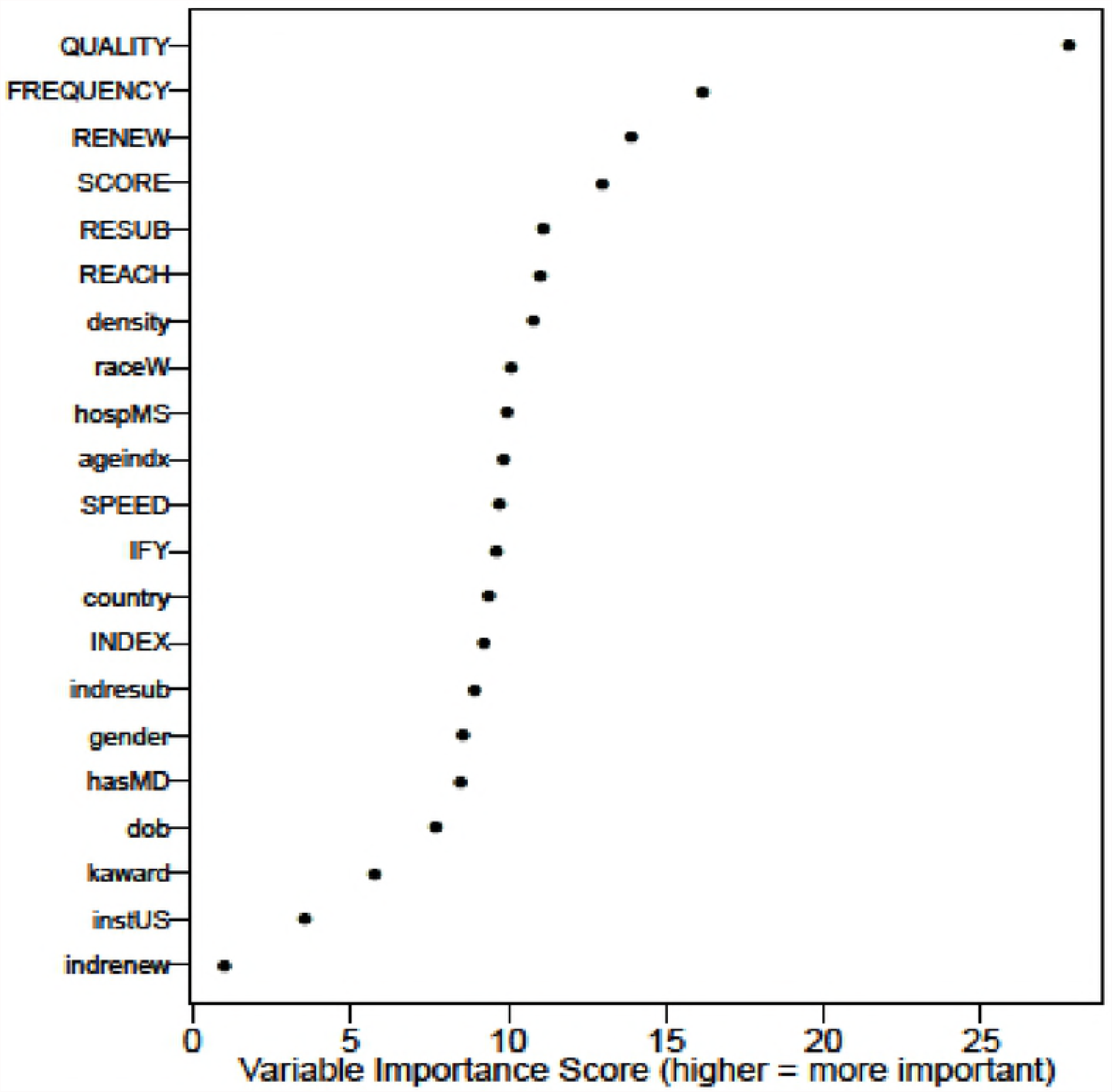
ENI Index Award Grant Activity Types

The vast majority of index awards (96%) were new (Type 1) awards, while a small proportion (4%) were renewal (Type 2) awards, meaning another PI began the project, but the study ENI submitted the competing renewal application (Table 6). Slightly more than half of the index awards (53%) were from resubmission applications, i.e. applications that had been revised from prior unfunded versions (formerly called amended applications). A small proportion of index awards (7%) were sub-projects. That is, the ENI was a project director of a sub-project on a multi-project grant. The median percentile score of all index awards was 13.7 (mean 15.7).

**Table 6.**
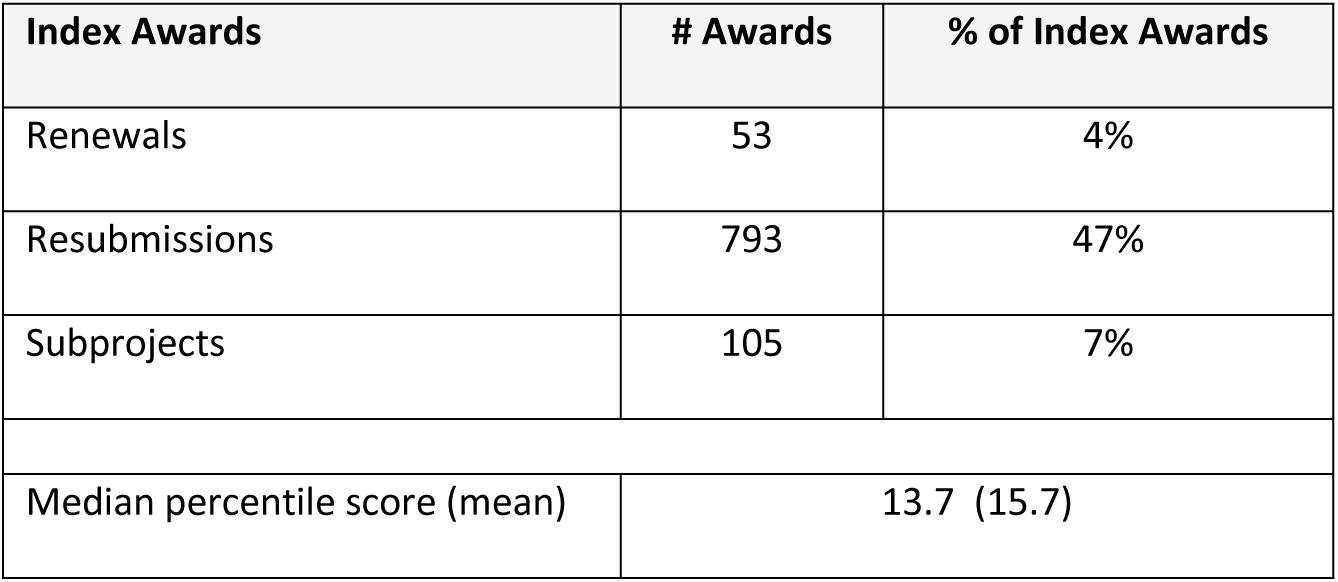
Index Award Characteristics.

| Index Awards | # Awards | % of Index Awards |
| --- | --- | --- |
| Renewals | 53 | 4% |
| Resubmissions | 793 | 47% |
| Subprojects | 105 | 7% |
| Median percentile score (mean) | 13.7 (15.7) |  |

### ENI Funding Outcomes

Our primary outcome of interest was whether an ENI received at least one new or renewal R01-e NIH grant award after the IFY. We refer to this outcome interchangeably as an “ENI funding outcome”, “ENI funding success”, or “ENI funding rate”. The ENI funding rate is the percentage of ENI within a particular comparison group, either across the entire period of the study or a specified period of time, successful in obtaining at least one R01-e grant after the IFY; it is calculated by dividing the number of ENI who received a post-IFY R01-e grant by the number of ENI in the comparison group or category.

Funding outcomes according to demographic, PI background, and index institutional characteristics were derived from the whole cohort (n = 1,496 ENI). Funding outcomes according to the PI’s application submission behaviors and application quality indices (i.e. all PI SCORECARD items) were derived using only ENI who submitted applications after the IFY (n = 1,322 ENI).

Ultimately, 57% of the cohort were funded and 43% were unfunded (Table 7). However, ENI from the first 4 cohort years had a statistically higher overall funding rate than ENI from the latter 4 years (60% versus 53%, respectively, *p* < 0.004). About 12% (174 ENI) did not apply for additional grants post-IFY. Of those who continued to apply, 65% were ultimately funded and 35% were not funded again. When we looked at the percentage of the cohort who remained active at least 5 years after receiving their index award – whether or not they had obtained new funding by then – 77% of the cohort remained, and 23% had dropped out (Table 8). Almost half (45%) of the unfunded ENI dropped out by their fifth year.

**Table 7.** ENI Funding Outcomes per Cohort Year.

| Cohort Year | # ENI in Cohort Year | # Unfunded ENI | # Funded ENI | % ENI Funded | % ENI Funded |
| --- | --- | --- | --- | --- | --- |
| 2003 | 236 | 79 | 157 | 66% | **60%** |
| 2004 | 185 | 73 | 112 | 61% |  |
| 2005 | 194 | 88 | 106 | 55% |  |
| 2006 | 167 | 67 | 100 | 60% |  |
| 2007 | 199 | 96 | 103 | 52% | **53%** |
| 2008 | 151 | 66 | 85 | 57% |  |
| 2009 | 197 | 98 | 99 | 50% |  |
| 2010 | 167 | 74 | 93 | 56% |  |
| All Years | 1496 | 641 | 855 | **57%** | ***p* < 0.004 \*** |

Numbers in table Include full cohort (n = 1,496 ENI)
\*Pearson's $\chi^2$ test, 2003-2006 versus 2007-2010, $\chi^2 = 8.3154$ , df = 1, *p*-value = 0.003931

**Table 8.** Percentage of ENI Who Remained in R01-e Applicant Pool 5 or More Years After Index Award.

| Outcome Group | # ENI Starting | # ENI Remaining | % ENI Remaining | % ENI Dropped out |
| --- | --- | --- | --- | --- |
| Unfunded | 641 | 353 | 55% | 45% |
| Funded | 855 | 800 | 94% | 6% |
| **Total** | **1496** | **1153** | **77%** | **23%** |

### Demographic, PI Background and Index Institution Characteristics

We found statistically significant differences between the funded and unfunded ENI according to some of the demographic characteristics (Table 9). Funded ENI were, on average, 3 years younger than unfunded ENI when they received their index award (median 40 y versus 43 y, *p* < 0.0001). In addition, funded ENI were born, on average, 2.5 years (median 2.2 years) later than unfunded ENI (*p <* 0.0001). There were no differences in percentages of males and females funded. A larger proportion of US-born ENI were funded compared to foreign-born ENI (63% versus 54%, *p =* 0.006). In terms of race and ethnicity, White ENI had the highest funding rate (61%), followed by AA/MR (59%), Asian (55%), and Hispanic (53%) ENI. Among the ENI for whom race/ethnicity data were missing or withheld (M/W), 34% were funded. There were 2 Native ENI in the cohort, and both were funded. When we performed a χ^2^ test for differences in funding rates across the race/ethnic groups – excluding the M/W – the differences were not statistically significant.

**Table 9.**
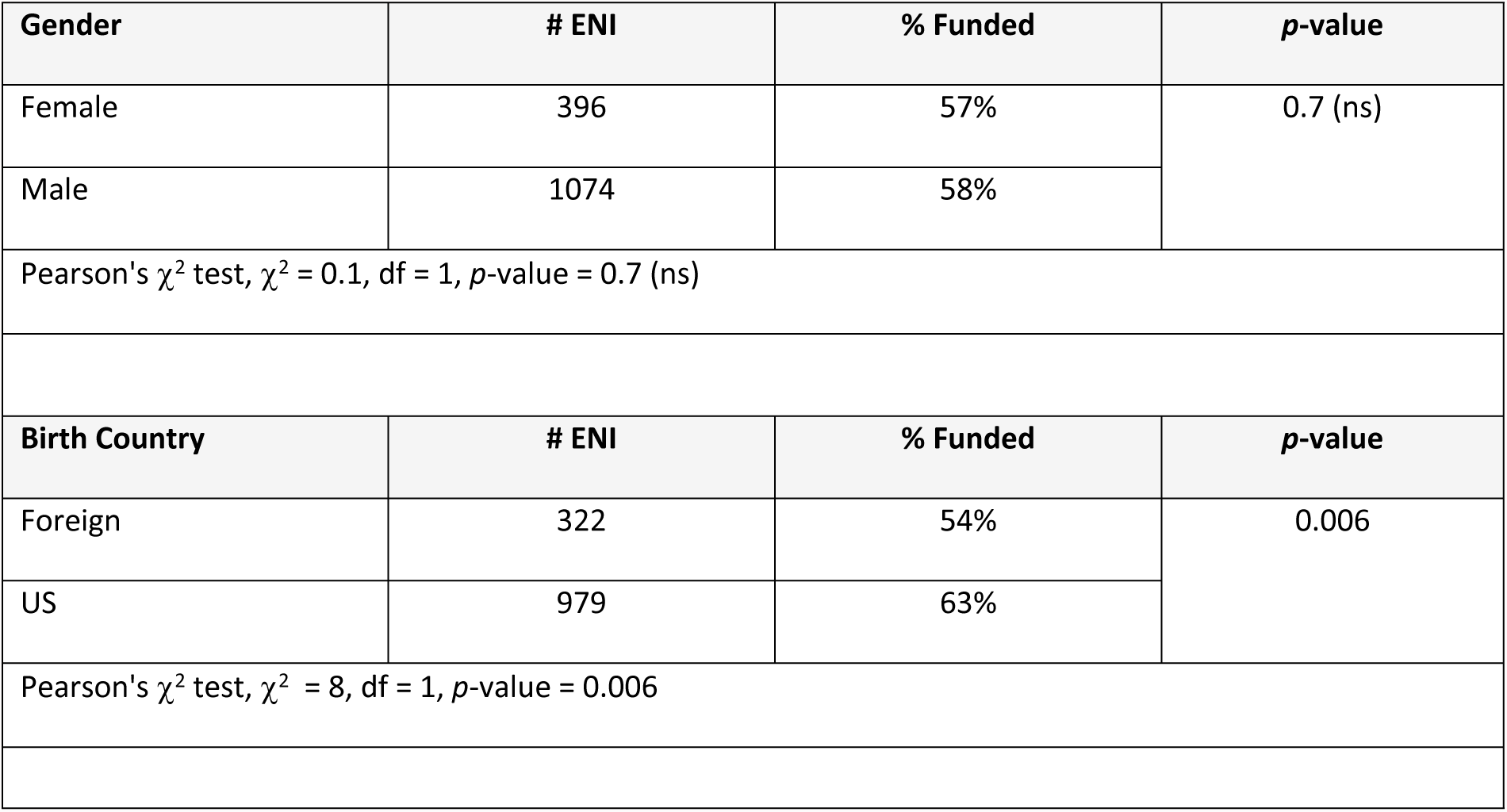
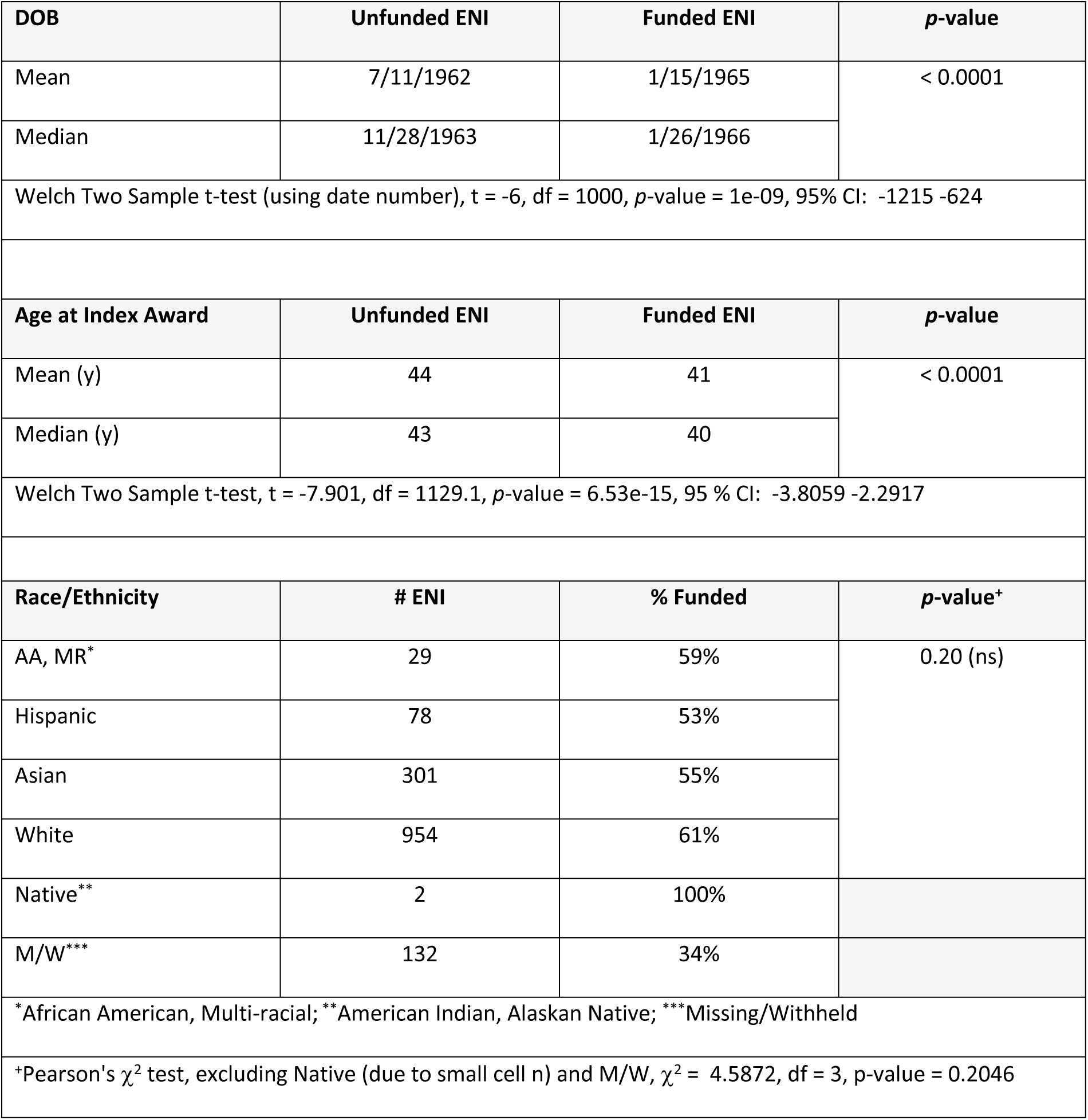
Funding Outcomes According to ENI Demographic Characteristics.

| Gender | # ENI | % Funded | *p*-value |
| --- | --- | --- | --- |
| Female | 396 | 57% | 0.7 (ns) |
| Male | 1074 | 58% |  |
| Pearson's $\chi^2$ test, $\chi^2 = 0.1$, df = 1, *p*-value = 0.7 (ns) | | | |
| Birth Country | # ENI | % Funded | *p*-value |
| Foreign | 322 | 54% | 0.006 |
| US | 979 | 63% |  |
| Pearson's $\chi^2$ test, $\chi^2 = 8$, df = 1, *p*-value = 0.006 | | | |

| DOB | Unfunded ENI | Funded ENI | p-value |
| --- | --- | --- | --- |
| Mean | 7/11/1962 | 1/15/1965 | < 0.0001 |
| Median | 11/28/1963 | 1/26/1966 |  |
| Welch Two Sample t-test (using date number), t = -6, df = 1000, p-value = 1e-09, 95% CI: -1215 -624 |  |  |  |
| Age at Index Award | Unfunded ENI | Funded ENI | p-value |
| Mean (y) | 44 | 41 | < 0.0001 |
| Median (y) | 43 | 40 |  |
| Welch Two Sample t-test, t = -7.901, df = 1129.1, p-value = 6.53e-15, 95 % CI: -3.8059 -2.2917 |  |  |  |
| Race/Ethnicity | # ENI | % Funded | p-value* |
| AA, MR* | 29 | 59% | 0.20 (ns) |
| Hispanic | 78 | 53% |  |
| Asian | 301 | 55% |  |
| White | 954 | 61% |  |
| Native** | 2 | 100% |  |
| M/W*** | 132 | 34% |  |
| *African American, Multi-racial; **American Indian, Alaskan Native; ***Missing/Withheld |  |  |  |
| *Pearson's $\chi^2$ test, excluding Native (due to small cell n) and M/W, $\chi^2$ = 4.5872, df = 3, p-value = 0.2046 | | | |

PI research training and institutional characteristics were all significantly associated with ENI funding outcomes (Table 10). ENI with an MD degree had a 64% funding rate, followed by those with an MD/PhD (62%), PhD (or equivalent) (55%), and Other degree (32%) (*p =* 0.004). Former recipients of an NIH K award had a higher funding rate than non-recipients (66% vs 55%, *p =* 0.002). ENI employed at a US institution at the time of their index awards had a higher funding rate than those employed at foreign institutions (60% vs 26%, *p <* 0.0001). Those whose index institutions were independent hospitals had the highest funding rates (68%), followed by medical schools (65%), other health or health related (e.g. not-for-profit, community service, international) organizations (64%), institutions of higher education (56%), and independent research organizations (52%) (*p* < 0.003).

**Table 10.** Funding Outcome According to PI Background and Index Institution.

| Degree Group | # ENI | % ENI Funded | *p*-value |
| --- | --- | --- | --- |
| MD | 289 | 63% | 0.04 |
| MD/PhD | 171 | 62% |  |
| PhD | 1016 | 55% |  |
| Other | 20 | 32% |  |
| Pearson's $\chi^2$ test, excluding Other, $\chi^2 = 6.5$, df = 2, p-value = 0.04 | | | |
| K Award | # ENI | % ENI Funded | *p*-value |
| Yes | 247 | 66% | 0.002 |
| No | 1249 | 55% |  |
| Pearson's $\chi^2$ test, $\chi^2 = 10$, df = 1, *p*-value = 0.002 | | | |
| Institution US | # ENI | % ENI Funded | *p*-value |
| US | 1371 | 60% | < 0.0001 |
| Foreign | 125 | 26% |  |
| Pearson's $\chi^2$ test, $\chi^2 = 50$, df = 1, *p*-value = 8e-13 | | | |
| Institution Type | # ENI | % ENI Funded | *p*-value |
| Independent Hospital | 130 | 68% | < 0.003 |
| Oth Health, Health-rel.,<br>community srvc. | 36 | 64% |  |
| Medical School | 429 | 64% |  |
| Institution of Higher Ed. | 636 | 56% |  |
| Research Organization | 156 | 52% |  |
| Pearson's $\chi^2$ test, $\chi^2 = 16.055$, df = 4, $p$-value = 0.002947 | | | |
| **ENI Density Tertile\*** | **# ENI** | **% ENI Funded** | **$p$-value** |
| 1 – High | 406 | 70% | < 0.0001 |
| 2 – Med | 454 | 65% |  |
| 3 – Low | 511 | 47% |  |
| Pearson's $\chi^2$ test, $\chi^2 = 60$, df = 2, $p$-value = 7e-13 | | | |

<sup>+</sup>High = 14 to 37 ENI per inst.; Medium = 6 to 13 ENI per inst.; Low = 1 to 5 ENI per inst.

Institutional ENI density was also significant, with ENI from high- and medium-ENI density institutions being more likely to be funded (70% and 65%, respectively) than ENI from low-ENI density institutions (47%) (*p <* 0.0001). Perhaps not surprisingly, given that institutions in the top 2 tertiles correspond to the top funded NIH institutions, the highest rate of ENI funding success (70%) was in the institution density tertile with the smallest number (21) of institutions (Fig 3).

**Fig 3.**
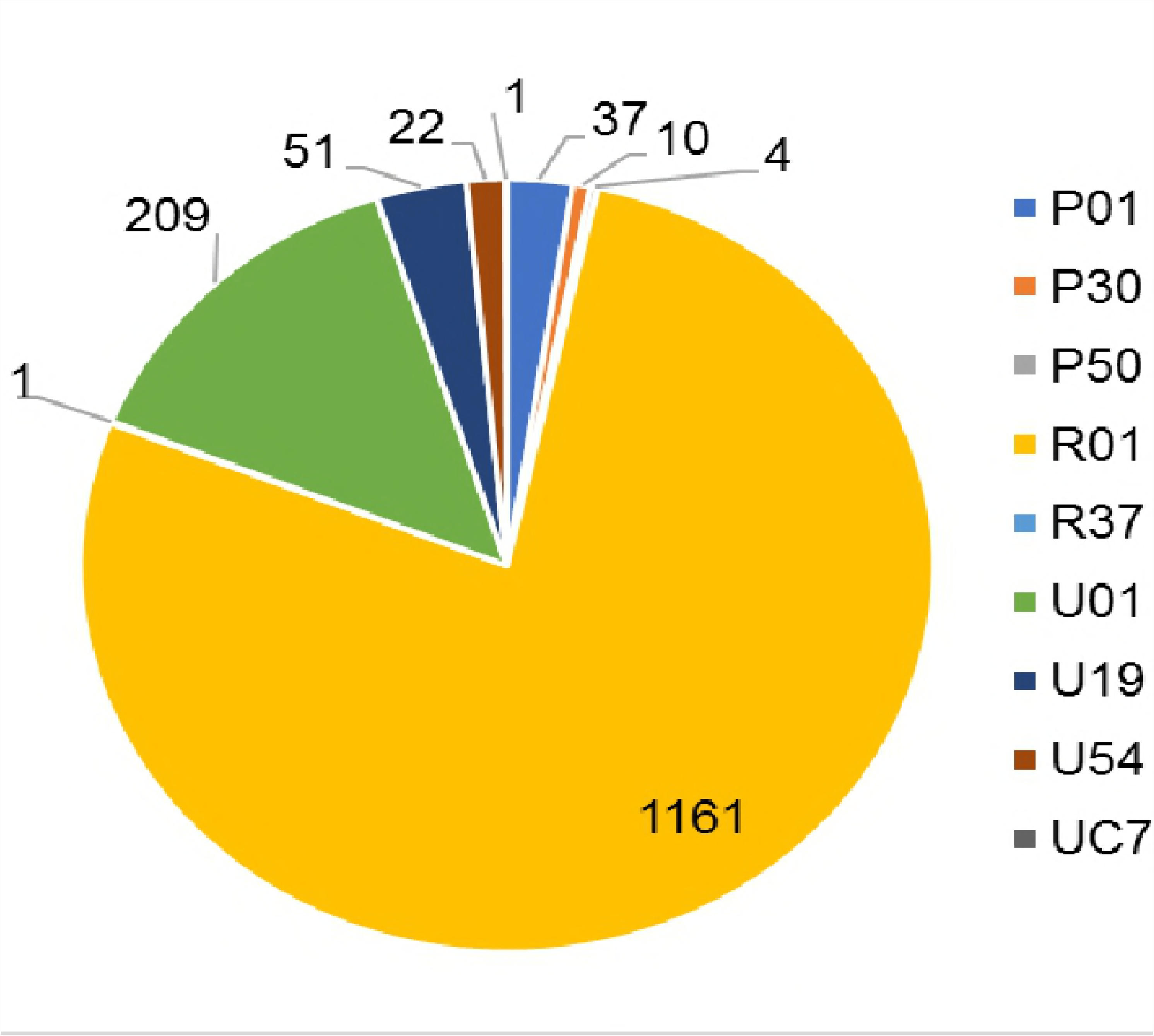
ENI Funding Outcomes by Index Institution ENI Density. Index institution density tertiles: 1 = 14 to 37 ENI per institution; 2 = 6 to 13 ENI per institution; 3 = 1 to 5 ENI per institution. ENI from institutions in tertiles 1 and 2 were more likely to be funded (70% and 65%, respectively) than ENI from institutions in tertile 3 (47%) (*p <* 0.0001). The highest ENI funding rate (70%) was in the 1^st^ tertile, which included just 21 institutions.

### PI SCORECARD Factors

There were several statistically significant differences between funded and unfunded ENI according to PI SCORECARD factors (Table 11). Funded ENI had: 1) lower median PI SCORES (22.4 versus 26.7, unfunded ENI, *p <* 0.0001); 2) lower median PI QUALITY (33% of applications triaged versus 50% triaged, unfunded ENI, *p* < 0.0001); 3) lower median INDEX scores (12.4 versus 14.9, unfunded ENI, *p* = 0.005); 4) higher median FREQUENCY rates (1.1 applications per year versus 0.8 applications per year for unfunded ENI, a difference of 27%, *p* = 0.0004); 5) faster median SPEED from IFY to next application (0.7 years versus 1.3 years, unfunded ENI, *p* = 0.005); 6) lower median REACH percentages (86% of applications to a single IC versus 100% for unfunded ENI, *p* = 0.004); 7) greater median RENEW percentages (17% of applications as renewals versus 10% as renewals, unfunded ENI, *p* < 0.0001); and 8) longer median ACTIVE times (8.7 years from IFY to final grant application (or FY 2016), versus 5.4 years for unfunded ENI, *p* < 0.0001). There was no difference between funded and unfunded ENI in the median percentage of the PI’s applications submitted as resubmissions (RESUB), with 30% for funded ENI and 33% for unfunded ENI (*p* = 0.30, n.s.).

**Table 11.** Funding Outcomes According to PI SCORECARD Factors.

| Factor | Unfunded ENI* | Funded ENI* | Test | *p*-values |
| --- | --- | --- | --- | --- |
| SCORE | 26.7 pctl | 22.4 pctl | Welch Two Sample t-test, $t = -2.8$, $p$-value = 0.005, 95 % CI: -2.886 -0.520 | < 0.005 |
| QUALITY | 50% | 33% | Welch Two Sample t-test, $t = -16$, $p$-value <2e-16, 95 % CI: -0.2174 -0.1698 | < 0.0001 |
| **INDEX** | 14.9 pctl | 12.4 pctl | Welch Two Sample t-test, $t = -6.0015$, $p$-value = $2.591\text{e-}09$, 95 % CI: -4.7354 -2.4021 | < 0.0001 |
| **FREQUENCY** | 0.8 apps/y | 1.1 apps/y | Welch Two Sample t-test, $t = 3.9$, $p$-value = $1\text{e-}04$, 95% CI: 0.0836 0.2549 | < 0.0001 |
| **SPEED** | 1.3 y | 0.7 y | Welch Two Sample t-test, $t = -5.3$, $p$-value = $1\text{e-}07$, 95% CI: -0.9144 -0.4221 | < 0.0001 |
| **REACH** | 100% | 86% | Welch Two Sample t-test, $t = -2.9$, $p$-value = 0.004, 95% CI: -0.0733 -0.0143 | 0.004 |
| **RENEW** | 10% | 17% | Welch Two Sample t-test, $t = 4.3$, $p$-value = $2\text{e-}05$, 95% CI: 0.0274 0.0734 | < 0.0001 |
| **RESUB** | 33% | 30% | Welch Two Sample t-test, $t = -1.1$, $p$-value = 0.3, 95% CI: -0.0351 0.0094 | 0.3 (ns) |
| **ACTIVE** | 5.4 y | 8.7 y | Welch Two Sample t-test, $t = 22.088$, $p$-value < $2.2\text{e-}16$, 95% CI: 3.2698 3.9073 | < 0.0001 |

\*Medians displayed

Given that ENI entered the cohort at different times, and funded ENI were generally born later than unfunded ENI, we further investigated some of the significant SCORECARD factors, to rule out or adjust for age and time effects. We repeated the analysis of ACTIVE for each cohort year. There were significant differences in the lengths of time ENI were ACTIVE between funded and unfunded ENI in every cohort year, except in 2010 (S1 Table.) The largest difference was in 2003 (6.3 y), and differences gradually diminished with each subsequent cohort year. This would be expected, as proportionally more of the unfunded ENI from the early years would have dropped out, and proportionally more of the funded ENI would have remained active.

Because funded ENI were on average 2.5 years younger than unfunded ENI, we wanted to be sure the difference in FREQUENCY of application submission per PI was not the result of their age difference. We took all ENI applications (approximately 10,000 after excluding subprojects) and plotted the number of applications submitted against PI age at time of submission (S1 Fig). The plot showed that even though funded ENI submitted many more applications than unfunded ENI (as expected), there was an identical pattern of application submission frequency relative to PI age at time of submission in both groups. ENI submitted the most applications between the ages of 42 and 44 years, very few applications before the age of 35, and they continued to submit applications through their mid-sixties, in both groups. These findings confirm that age *per se* was not driving the higher frequency of application submission among the funded ENI.

We also examined whether PI SCORECARD findings held up across cohort years and study observation years. First, we looked at index award scores (INDEX) according to ENI cohort year (Fig 4). Index scores did vary from year to year, but that was not surprising because NIAID’s R01 payline does change from year to year [42]. However, unexpectedly, index award scores of funded and unfunded ENI were statistically different in cohort years 2003, 2005, 2006, 2007, and 2009. In 2004, 2008 and 2010, index scores were not statistically different. Two factors come into play in understanding this. First, a normal part of the NIAID funding process every year is to select some additional R01 applications that did not score within the payline, and award funding (a process called “select pay”) [43]. Priority for select pay is frequently given to new investigators (NI). Second, starting in 2006, NIAID established special (preferential) paylines for NI R01 grants. So, between 2006 and 2010, there were NI preferential paylines, in addition to select pay, which were used to fund R01 grants *above the NI payline*. We looked at ENI index award scores between FYs 2006 and 2010, and found that ENI whose awards were paid at or below the NI payline were more often successfully funded, than those whose awards were paid above the NI payline, and the difference was statistically significant (60% versus 48%, χ^2^ test, *p-*value < 0.004, S2 Table). Thus, some NI received their index awards having scores well above normal paylines and this may have conferred a disadvantage later in competing effectively at normal paylines.

**Fig 4.**
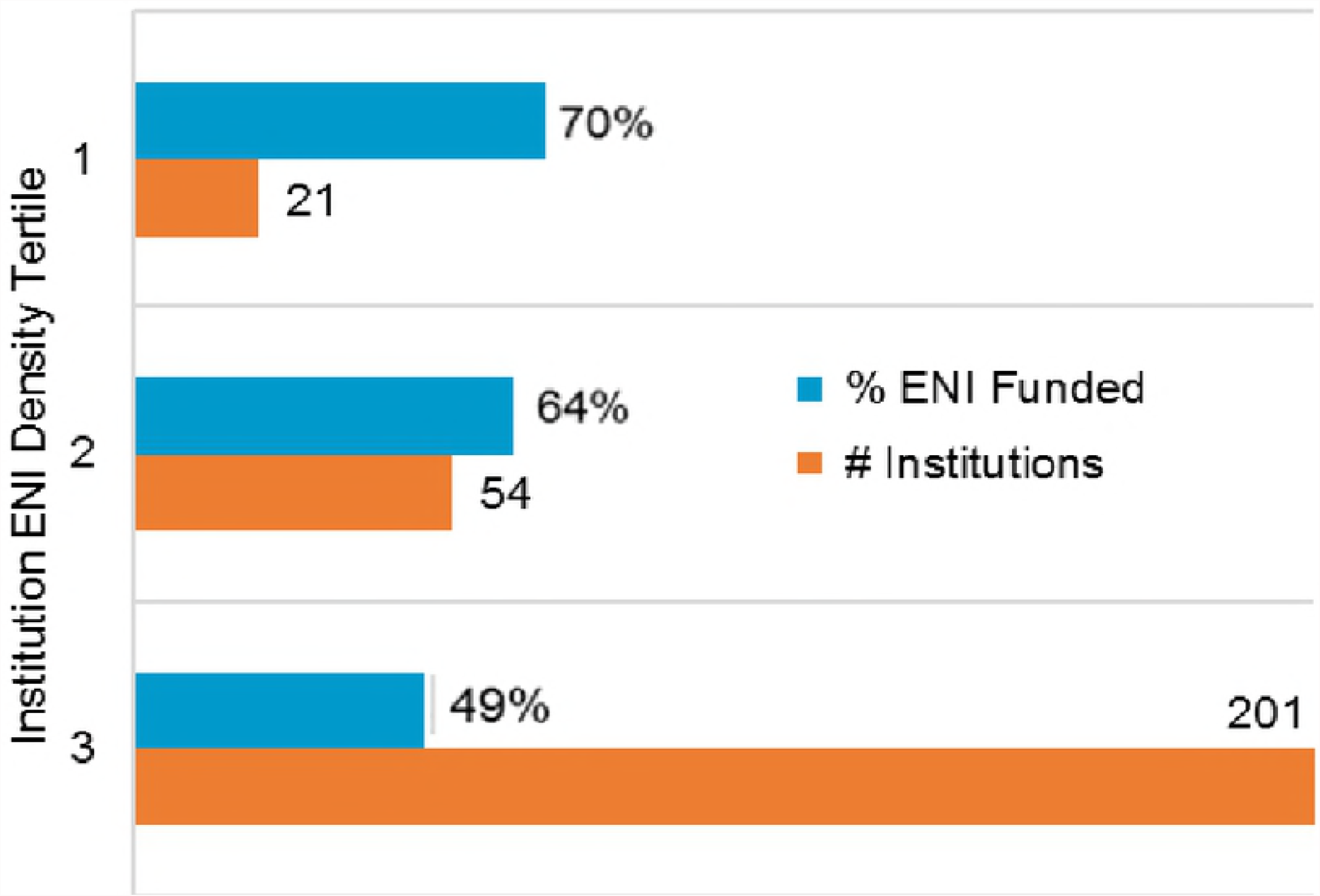
ENI Index Award Scores by Cohort Year. Index scores varied from cohort year to cohort year, a result of normal NIAID R01 payline changes from year to year. Unexpectedly, index award scores of funded and unfunded ENI were statistically different in cohort years 2003, 2005, 2006, 2007, and 2009. In 2004, 2008 and 2010, index scores were not statistically different.

Next, we examined the year-over-year differences between funded and unfunded ENI in terms of the total number of applications they submitted and how many of them were triaged (versus scored, Fig 5). For this analysis we included only applications from the 1,322 ENI who applied for grants post-IFY, and we excluded index awards. Between FYs 2003 and 2016, funded ENI submitted a total of 8,026 applications, of which 3,292 (39%) were triaged; unfunded ENI submitted a total of 2,202 applications, of which 1,470 (63%) were triaged. In both groups, the number of applications submitted continued to increase from 2003 through 2010, while new ENI were still coming into the cohort (black dotted line in Fig 5). After 2010, the number of applications per year submitted by unfunded ENI remained relatively steady through 2016, as did the proportion of those applications triaged. In the funded group, the number of applications submitted each year continued to increase after 2010, but at a less steady pace than 2003 to 2010, and the proportion of applications triaged each year was relatively constant. Thus, funded ENI not only submitted more applications per year than unfunded ENI, even while the cohort was still growing, but consistently had a higher proportion of their applications scored, rather than triaged. The differences we see in years 2003 through 2010 suggests funded ENI had an ability *from the start* to write higher quality applications than unfunded ENI. This superior grant writing ability appears to be another early advantage funded ENI had, which may have conferred a lasting benefit to them.

**Fig 5.**
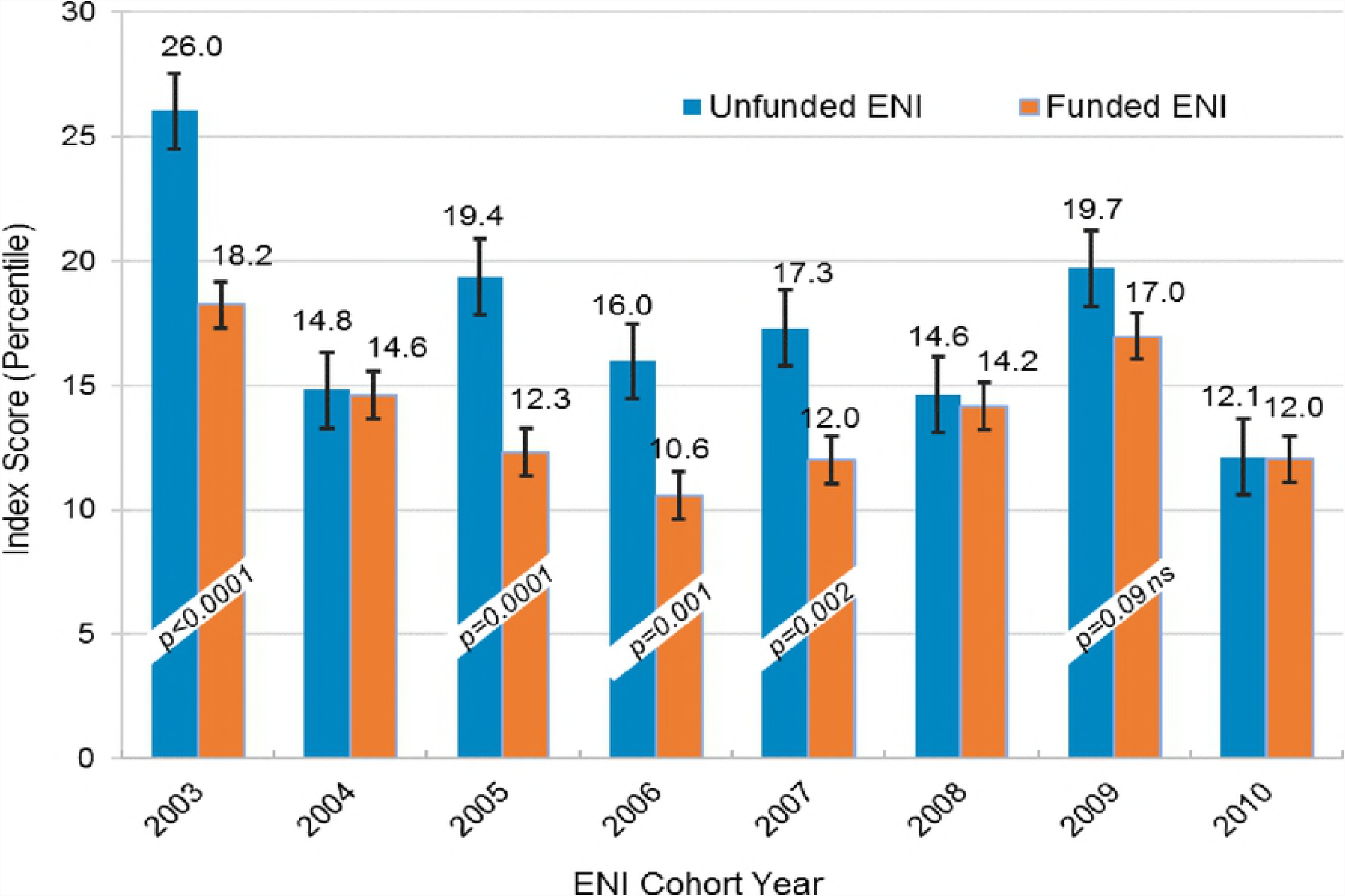
Applications Scored and Triaged from Funded and Unfunded ENI. Figure includes 10,228 applications from 1,322 ENI who submitted applications after the IFY. (Index applications are excluded.) Funded ENI submitted 8,026 applications, of which 39% were triaged; unfunded ENI submitted 2,202 applications, of which 63% were triaged. The number of applications from both groups increased between 2003 and 2010, while the cohort was still growing (black dotted line), but more rapidly from funded ENI. Funded ENI consistently had fewer of their applications triaged, even in the early years, suggesting they had an early advantage in grant writing ability.

Lastly, we scrutinized the difference between funded and unfunded ENI in terms of average PI application scores over time, starting in 2011 when no additional ENI were entering the cohort. That is, we used all scored applications submitted by ENI who continued to apply for grants between FYs 2011 and 2016 (n = 3,093 applications total). For each year, an average *PI ANNUAL Score* was calculated (for both funded and unfunded ENI) as follows: any ENI who submitted one or more scored applications in the year received an individual *ANNUAL Score*, equal to the average percentile score of his/her scored applications. If an ENI submitted only one scored applicaton, his/her individual ANNUAL Score was equal to the score of that application. Each year’s PI ANNUAL Score was the average of the individual ANNUAL Scores – that is, the sum of the individual ANNUAL Scores, divided by the number of individual ANNUAL Scores. As such, only ENI who submitted scored applications in a given year contributed to that year’s average PI ANNUAL Score. As shown in Fig 6, average PI ANNUAL Scores were markedly different between the funded and unfunded ENI, with funded ENI having PI ANNUAL Scores about 10 percentile points lower each year between 2011 and 2016. Overall, the mean PI ANNUAL Score for the funded ENI was 10.9 percentile points lower than that for the unfunded ENI (23.9 versus 34.8, respectively, *p* < 0.0001).

**Fig 6.**
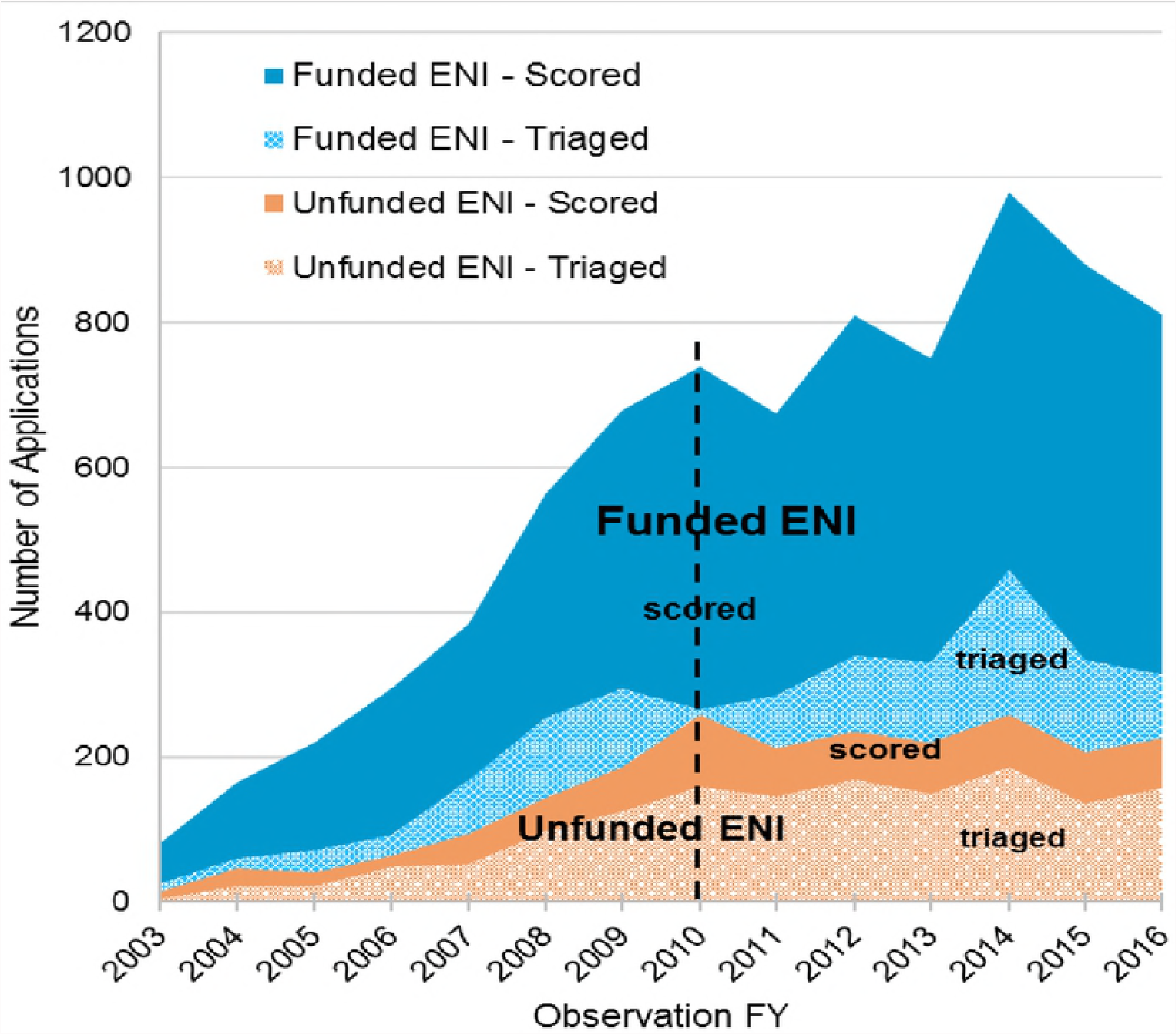
Average PI ANNUAL Scores, FY 2011 – FY 2016. The PI ANNUAL Score each year is the average of individual PI ANNUAL scores of ENI who submitted scored applications. Average PI ANNUAL Scores were markedly different between the funded and unfunded ENI, with funded ENI having PI ANNUAL Scores about 10 percentile points lower each year between 2011 and 2016. (Welch Two Sample t-tests: in all years *p*-values < 0.0001.)

When we looked at the distributions of PI average application ANNUAL Scores within the two ENI groups, in each year, funded ENI not only had a broader range of ANNUAL Scores than unfunded ENI, they also submitted many more scored applications than unfunded ENI (S2a and S2b Figs). S2a Fig shows the distribution of ANNUAL Scores for funded and unfunded ENI in FY 2012; S2b Fig shows the distribution of scores as well as the cumulative numbers of ENI contributing to those scores in each group. FY 2012 is typical of all the years between 2011 and 2016.

### Regression Analyses

Having identified numerous statistically significant associations between ENI funding outcomes and independent demographic, PI background, institutional, and PI SCORECARD variables, we wanted to understand the strength of each of these variables in predicting, individually and collectively, the likelihood of ENI funding success. We performed univariate and multivariate logistic regression analyses, and discuss our results.

The strength of individual independent variables in predicting ENI funding success was assessed using univariate logistic regression analyses. Each of the independent variables was converted to a binomial, with values 1 or 0 indicating the test condition was met or not met, respectively. The results are shown in Table 12. All but 3 of the 21 variables tested were statistically significant predictors. The strongest predictors were among the PI SCORECARD factors and included the PI submitting: more renewal applications (RENEW); more applications to different NIH ICs (REACH); more applications per year (FREQUENCY); and fewer applications triaged (QUALITY). Having a lower index award score (INDEX) was also predictive of funding success. RENEW, REACH and FREQUENCY increased the odds of an ENI being funded by 2.8-, 2.8-, and 2.4-fold, respectively. QUALITY and INDEX each increased the odds by 1.6-fold. Demographic and institutional factors that were also highly predictive included the PI being: younger at receipt of the index award; younger generally (i.e. born later); white; and employed at the time of the index award at a US institution, an independent hospital or medical school, or an ENI-dense institution. Institutional EEI density – which correlates with NIH funding level – increased the odds of an ENI being funded by almost 3-fold. Additional PI SCORECARD factors that increased the chance of funding success by about 30% each included the PI having a lower average application score (SCORE), and a shorter time between the index award and the next application (SPEED). Other demographic and institutional factors predictive of funding included the PI having: an MD or MD/PhD degree, a prior K award, the index award before 2006; a renewal index award; and US birth. Having more resubmission applications (RESUB), a resubmission index award, and gender, were not statistically significant predictors.

**Table 12.** Univariate Regression of Independent Variables on ENI Funding Success.

| Predictor Variable* | Predictor Value Tested<sup>+</sup> | Odds Ratio<sup>o</sup> | *p*-value |
| --- | --- | --- | --- |
| **RENEW** | % PI's apps=renewal ≥ median | 2.83 | < 0.0001 |
| **REACH** | % PI's apps to single NIH IC < median | 2.75 | < 0.0001 |
| **FREQUENCY** | PI's average # of apps per year ≥ median | 2.44 | < 0.0001 |
| **QUALITY** | % PI's apps triaged < median | 1.63 | < 0.0001 |
| **INDEX** | PI's index award score < median | 1.63 | < 0.0001 |
| **Ageindx** | Age at index award < median | 2.10 | < 0.0001 |
| **density** | Index inst density = 1 or 2 | 2.73 | < 0.0001 |
| **hospMS** | Index inst type = independent hospital or medical school | 1.76 | < 0.0001 |
| **instUS** | Index inst = US inst | 4.17 | < 0.0001 |
| **dob** | Date of birth ≥ median (PI is younger) | 1.62 | < 0.0001 |
| **raceW** | Race = white | 1.55 | < 0.0001 |
| **kaward** | Had K award | 1.60 | < 0.005 |
| **indrenew** | Index app = renewal | 2.64 | < 0.005 |
| **Birth** | Birth country = US | 1.44 | < 0.005 |
| **SCORE** | PI's average app score < median | 1.33 | < 0.05 |
| **SPEED** | Time between index award and first<br>subsequent app < median | 1.27 | < 0.05 |
| **hasMD** | Degree = MD or MD/PhD | 1.37 | < 0.05 |
| **IFY** | IFY < median (< 2006) | 1.31 | < 0.05 |
| **RESUB** | % PI's apps=resub ≥ median | 1.09 | ns |
| **gender** | Gender = male | 1.05 | ns |
| **indresub** | Index = resub | 1.01 | ns |

\*PI SCORECARD variables in all CAPS; \*app=application, inst=institution, resub=resubmission; ° increase in odds of being funded with factor.

In addition to wanting to know the impact of individual independent variables on ENI funding success, we wanted to understand how the effect of these variables changed when considered in a multivariate model. When we fit all the independent variables into a generalized linear model using multvariate logistic regression, we found that QUALITY, SCORE, RENEW, and FREQUENCY prevailed as the most highly significant predictor variables, with QUALITY conferring a 5-fold increase in the odds of ENI funding success and SCORE, RENEW and FREQUENCY each conferring more than a 2-fold increase in the odds of funding success (S3 Table). RESUB, REACH, density, and ageindx also remained statistically significant predictors, each conferring, on average, a 1.6-fold increase in the odds of success. Most of the other variables – primarily demographic and institutional factors – lost their statistical significance or remained only weakly significant in the multivariate model.

Finally, we wanted to better understand how important each of the predictive factors were relative to one another when considered altogether. For this analysis, we used Random Forest Variable Importance (RFVI) modeling, which accounts for correlations and any interactions among the variables, and ranks the independent variables in order of their importance. By far, the percentage of the PI’s applications triaged (QUALITY) was the strongest predictor in the model (Fig 7). The next most important predictors (in order of their importance) were the PI’s: average number of applications submitted per year (FREQUENCY); percentage of renewal applications (RENEW); average application score (SCORE); percentages of resubmission applications (RESUB) and applications submitted to multiple NIH ICs (REACH); and the index institution ENI density. The variables with the least importance were having a Type 2 index award and a US index institution; having a K award ranked just slightly higher. All the remaining variables had approximately equivalent predictive strength.

**Fig 7.**
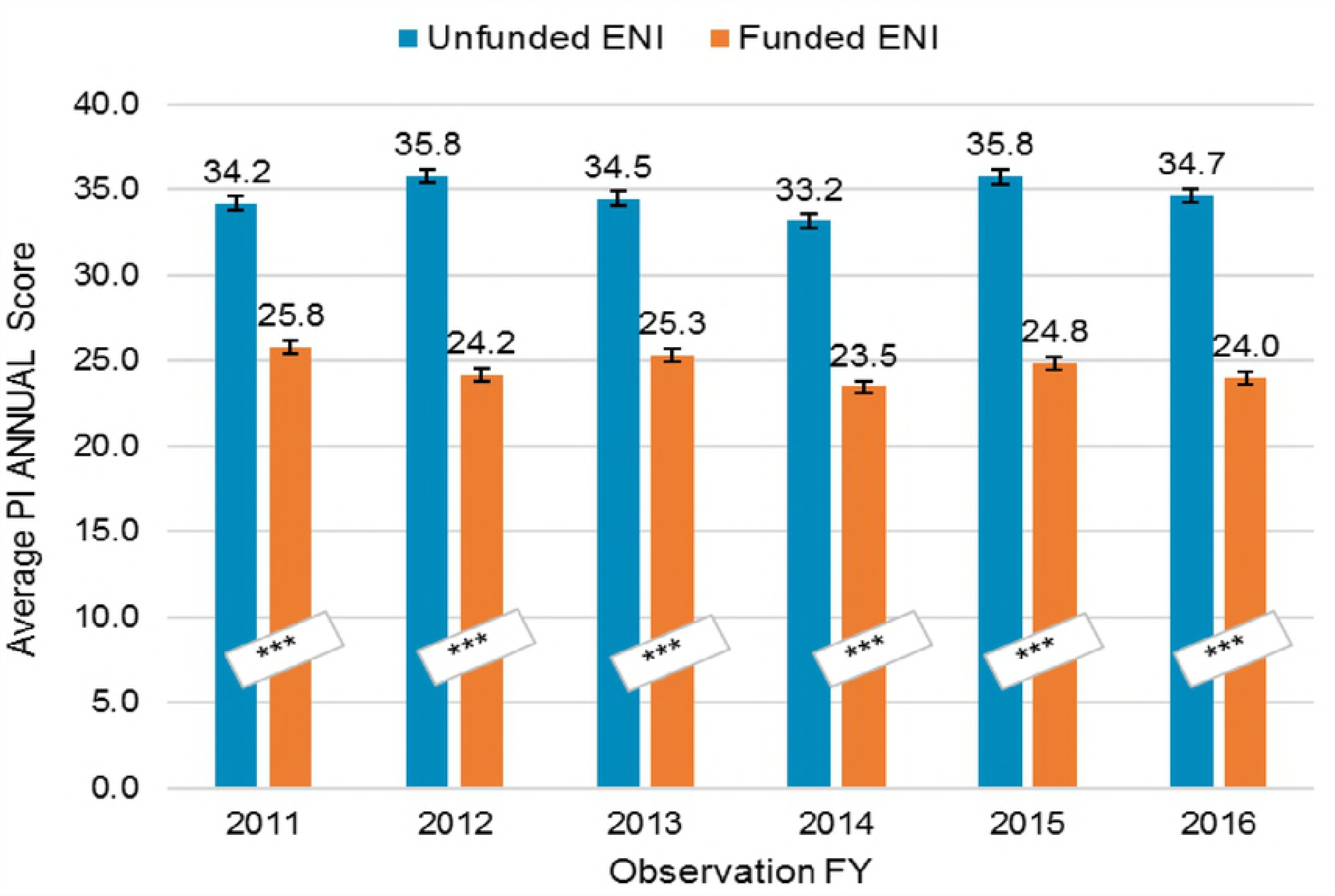
Variable Importance in Prediction of ENI Funding Success. RFVI analysis ranked all independent variables included in multivariate modeling in order of importance in predicting ENI funding success. The strongest predictor was the percent of the PI’s applications triaged (QUALITY), followed by the PI’s: average applications per year (FREQUENCY); percent of renewal applications (RENEW); average application score (SCORE); percent of resubmissions (RESUB) and applications to multiple NIH ICs (REACH); and the index institution ENI density. Having a K award, a renewal index award and a US index institution, were the least important predictors. All of the other variables had approximately equal predictive strength.

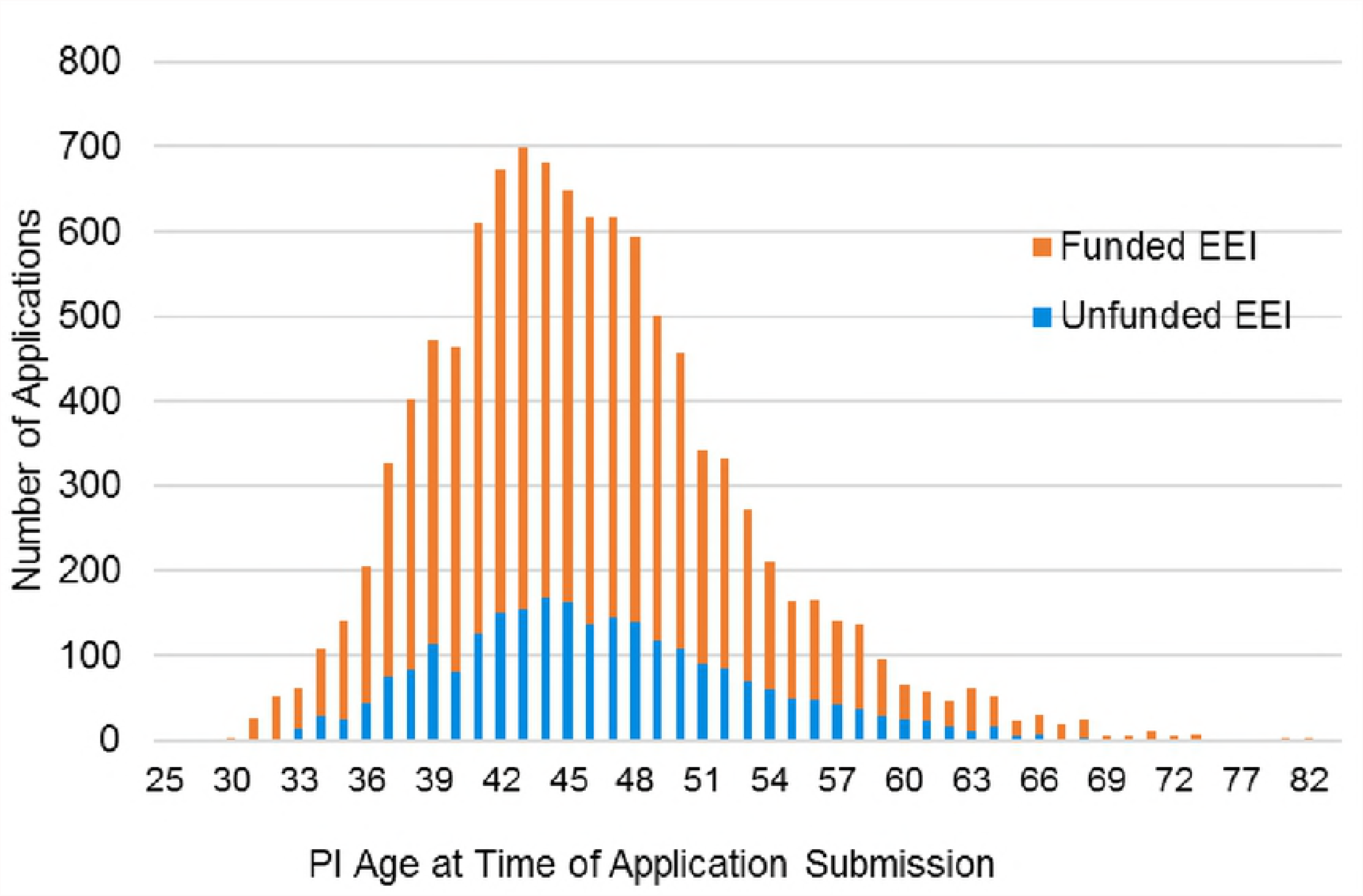

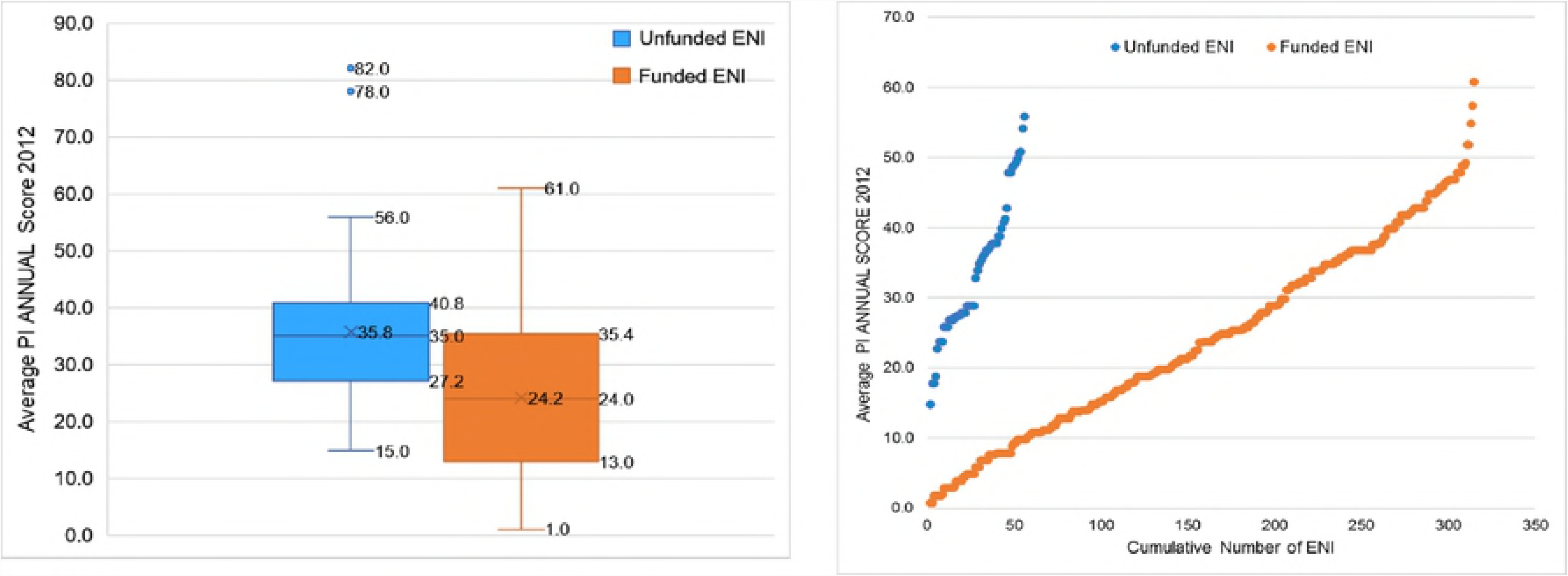

## Discussion

### The Out-of-Balance Biomedical Workforce

There is widespread recognition that the current funding structure of the US biomedical research enterprise is severely imbalanced [1, 3, 44-46]. Science organizations and thought leaders have called for broad structural reforms and proposed strategies for reversing these declines [4, 47-49]. Some of the many proposed solutions include: amplifying programs to support early- and mid-career stage investigators [6, 50-52]; funding people instead of projects [46, 53, 54]; reducing the size of laboratories, of awards, or of numbers of NIH grants an investigator may hold at any one time [6, 46, 47, 55, 56]; and reducing NIH support for investigator salaries and reliance on soft-money positions [2, 47].

These solutions have various levels of support in the biomedical research community, but they all represent significant structural and/or institutional reforms, and as such, none are easy to implement. That said, more practical answers may arise from a better understanding of how funding agencies and institutions can better support early-career scientists who are most at risk. This understanding could identify specific interventions by institutions and funding agencies, and behaviors of the researchers themselves, that could enhance their competitiveness.

### Factors Contributing to ENI Success in a Hypercompetitive Environment

We studied a cohort of 1,496 ENI who received their first R01-e awards from NIAID between FYs 2003 and 2010. This was a period of no overall growth in the NIH inflation-adjusted budget and the steepest declines in success rates in NIH history [57]. Ironically, the number of research grant applications continued to grow during this period, primarily due to an increase in the absolute number of applicants [58, 59]. We tracked the cohort’s ENI grant applications and funding outcomes, from their first R01-e awards through their final application submissions, or FY 2016, whichever came first. Despite the challenges facing these early-career scientists during this period, over half of the cohort was successful in obtaining subsequent NIH funding.

We were able to identify many factors that differentiated ENI who were successful in obtaining additional R01-e grants after their index award) from those who were not successful. Using these factors, we constructed a model that would predict the likelihood of a cohort ENI successfully obtaining additional funding. Characteristics that differentiated successful from unsuccessful ENI fell into 3 major categories: 1) unalterable PI personal attributes; 2) PI background and institutional factors; and 3) PI grant quality and grant submission behaviors.

ENI from the early cohort years (2003 – 2006) had higher funding rates than ENI from the later cohort years (2007 – 2010). This may be due in part to higher R01 paylines in the years the early cohort ENI competed for new funding. For most of the early cohort ENI it may also be due to an absence of preferential NI paylines: early cohort ENI had to compete against established investigators for their first R01-e awards without the benefit of preferential paylines. This may have prepared them better for competition later at normal paylines.

Funded ENI were, on average, 2.5 years younger than unfunded ENI, within each cohort year. Funded ENI also received their index awards at an average age of 40 years, compared to an average age of 43 years among unfunded ENI. We considered the possibility that the earlier age at index award may reflect, in part, changes in the NIH new investigator policies between 2007 and 2009, including establishment of numerical benchmarks for new investigator awards, comparable type-1 R01 success rates between new and established investigators, and identification of “Early Stage Investigators” [60]. In our study, ENI from cohort years 2003 through 2006 would not have benefited from these policies. We did not find significant differences between funded and unfunded ENI from the first 4 cohort years compared to those from the second 4 cohort years in birth date and age at index award. This suggests that any effect of age on subsequent ENI funding success was independent of NIH policy changes during the observational period. Why such seemingly small differences in age might make a difference in ENI outcomes remains unclear.

Our findings that ENI funding rates were highest in independent hospitals and medical schools, and that ENI with MD or MD/PhD degrees had higher funding rates than ENI with PhD degrees within independent hospitals, medical schools, and research organizations, is consistent with NIH reporting [61-63]. Historically, medical schools have received the largest share of NIH funding [40].

We also found higher ENI funding rates in institutions with the highest ENI densities and learned that our ENI density tertiles correspond well with institutional level of NIH funding. Ginther et al., in their study of over 40,000 investigators from FYs 2000 to 2006, reported that working at one of the top 30 institutions, ranked by total NIH grant funding, increased an investigator’s R01 award probability by 9.7 percentage points, and those working at institutions ranked 31-100 increased R01 award probability by 6.1 percentage points [64]. In our study, there were 75 institutions (27% of all the institutions) in the top two ENI density tertiles. The 860 ENI (i.e. 2/3 of the cohort) who received their index awards while at an institution in one of these two tertiles had an average funding rate of 67%. In contrast, 511 ENI employed at the time of their index award at institutions in the bottom tertile, comprised of 201 institutions (i.e. 73% of the institutions), had a funding rate of 47%. It is tempting to speculate that institutions with relatively high ENI densities provide an environment where younger early-career researchers can share ideas and pursue more innovative projects.

PI SCORECARD factors were surprisingly effective in identifying factors preferentially associated with funded ENI. Compared to unfunded ENI, funded ENI submitted 20% more applications per person per year, nearly 30% more of their applications to different NIH ICs, 30% more of their applications as renewals, and their first post-IFY applications on average about 8 months sooner. Submission of more applications to different NIH ICs suggests these projects had broad scope and relevance, opening extra opportunities to seek funding from multiple ICs. Submission of more renewal applications suggests these ENI had achieved most of the objectives of their original grants, leading them to be more strategic and competitive: NIH data show, for both new and experienced investigators, renewal applications have higher success rates than new applications [65]. But NIH data also show that first renewal applications from ENI have lower success rates than all renewal applications from established investigators, because all renewals from established investigators include new as well as long-term projects, which have even higher success rates than first renewals [66].

Finally, if we compare our study cohort with the 1983-2003 NIAID cohort depicted in Fig 1, we can make two observations: First, in our study, the steepest drop out occurred between the 4^th^ and 5^th^ year after the index award, similar to the earlier cohort. Yet, the median number of years our unfunded ENI remained active was 5.4, and 55% of the unfunded ENI were still active by 5 years, suggesting, perhaps, a slight lengthening of survival time for the more recent cohort. Second, over ¾ of our cohort (77%) remained active at least 5 years, compared to just 68% remaining after 5 years in the earlier cohort, again suggesting survival time may be improving. Additional years of follow-up of our study cohort will be needed before more definitive conclusions about change in ENI survival time can be made.

While relationships between submission behaviors and funding success most likely seems intuitive, or even predictable, we are unaware of other studies that have reported on these relationships quantitatively. A more nuanced observation from our study is that successful ENI displayed remarkable within-person consistency, not only in grant submission behavior, but across multiple behaviors associated with a higher odds likelihood of future funding.

We looked at the strength of the independent variables in predicting ENI funding success both the univariate and multivariate analyses revealed that the strongest, statistically significant predictor variables were: a low percentage of the PI’s applications triaged (QUALITY), frequent application submission by the PI (FREQUENCY), a low PI average application score (SCORE), submission of more renewal applications (RENEW) and more applications to different NIH ICs (REACH), a younger PI age at index award, and a high ENI-density index institution. Individually and collectively each of these factors conveyed 2- to 4-fold increases in the odds of an ENI being funded.

Finally, we used RFVI analysis to compute and rank all the predictor variables in terms of relative importance. Unlike the univariate and multivariate analyses, which identified and showed the impact of the predictor variables, the RFVI analysis revealed the order and relative importance of each one when all were included in the model. By far, the proportion of the PI’s applications triaged was the most important predictor, followed by the PI’s rate of application submission, application scores, and percentage of renewal applications. It is important to point out more effective grant writing and grant submission behaviors confer a strong cumulative advantage for ENI, especially when they submit high quality applications.

## Conclusion

Our study describes the characteristics of ENI from a NIAID cohort of first-time R01 investigators who were successful in obtaining new or renewal R01-e funding after their index award. They were successful despite a highly competitive funding environment that favored more senior investigators. Funded ENI began with a slightly better median index score than unfunded ENI (2.5 percentile points better), and an ability to write better applications (fewer were triaged) even while the cohort was still growing, and these characteristics may have conferred cumulative, lasting benefits to them. Clearly, the divergence between the two groups grew over time. When we compared grant submission behaviors and grant quality indices, what emerged was the profile of the tenacious, successful ENI, who developed superior grant writing skills, superior grant submission strategies, and projects with broad relevance and scope.

It should be noted that this study did not examine the potential role of the specific scientific areas that were pursued by the successful and unsuccessful ENI, and whether there were any differences or trends. Because the NIAID supports research across a broad range of scientific areas (basic immunology and microbiology, pathogenesis of infectious diseases, immune-mediated diseases and transplantation, as well as translational and clinical research in these areas), the cohort reflects the broad mandate of the Institute. That said, our data do indicate that PIs who had the ability to submit applications to more than one IC had an increased likelihood of being successful. This implies that sciences areas that are more amenable to cross-cutting and trans-disciplinary research may confer an advantage to ENI working in these areas. Future work is needed to explore these possibilities.

Whether the characteristics displayed by the successful ENI were the results of better mentorship, institutional training resources, access to institutional core facilities, an innate ability to persevere, or all the above, is something about which we can only speculate. These factors are particularly important because several are obvious points of intervention by institutions and funding agencies.

## Acknowledgements

We would like to thank Dr. Jason Liang of the Division of Clinical Research of the NIAID for his help with the regression and random forest statistical methods used in this manuscript. Dr. Liang wrote the original R code for these analyses and provided invaluable assistance in interpreting their results.

## Author Contributions

**Conceptualization:** MJF PAH

**Data Curation:** PAH

**Formal analysis:** PAH

**Investigation:** PAH

**Methodology:** PAH

**Resources:** PAH

**Supervision:** MFJ

**Validation:** PAH MJF

**Visualization:** PAH MJF

**Writing** – original draft: PAH

**Writing – review & editing:** MJF PAH

## Supporting information

**S1 Appendix.** Details of cohort selection and identification of project start dates in IMPAC II

**S1 Table.** Mean Number of Years ENI ACTIVE Applying for NIH Grants

**S1 Fig.** Frequency of application submission by PI age at time of submission

**S2a Fig.** ENI Average Application ANNUAL Scores, 2012

**S2b Fig.** ENI Average Application ANNUAL Scores and Cumulative Numbers of ENI, FY 2012

**S2 Table.** ENI Funding Success According to Index Award Score Above or Below NI Payline

**S3 Table.** Multivariate Logistic Regression Model, Predictive Strength of Independent Variables on the Odds of ENI Funding Success

